# Barcoded Competitive Clone-Initiating Cell (BC-CIC) Analysis Reveals Differences in Ovarian Cancer Cell Genotype and Niche Specific Clonal Fitness During Growth and Metastasis In Vivo

**DOI:** 10.1101/2021.04.08.439098

**Authors:** Syed Mohammed Musheer Aalam, Xiaojia Tang, Jianning Song, Jamie Bakkum-Gamez, Mark E. Sherman, Upasana Ray, Viji Shridhar, David J.H.F. Knapp, Krishna R. Kalari, Nagarajan Kannan

## Abstract

During oncogenesis, pathogenic clones develop which contain cells capable of spreading throughout the body, ultimately compromising vital organ functions and physiology. Understanding how metastatic clones develop and spread is critical for improving cancer treatments. However, our understanding of these processes has been hampered by a paucity of quantitative methodologies to comprehensively map, track and characterize such clones. To address this shortcoming, we have developed a DNA barcoding and next-generation sequencing based system-wide clonal tracking technology integrated with a computational data analysis pipeline called Clone-Initiating Cell (CIC) Calculator. The CIC Calculator interfaces with the CIC Morbus Mandala (CIC-MM) plot, a novel tool to visually comprehend and detect four distinct categories that explains their complex relationships with various tissues/organ sites. Further, we describe machine learning approaches to study CIC number, frequency, and estimate clone size and distribution demonstrating distinct growth patterns, and their inter-relationships and their routes of metastatic spread at clonal resolution. We demonstrate these methodologies, using our novel multifunctional lentiviral barcode libraries, and specifically barcoded tubal-ovarian metastatic OVCAR5 cell lines (engineered to express varying levels of metastasis promoting LRRC15 gene) and co-injected cells in a competitive CIC assay into tubal or ovarian sites in highly immunodeficient NSG mice. DNA was isolated from primary tumors, omental/bowel metastasis and system-wide anatomical site/organs. Amplicon sequencing libraries were constructed with spike-in-control barcodes (serving as internal calibration controls) to estimate absolute clone sizes. The computational pipeline CIC Calculator was then used to deconvolute and filter the data, set stringent thresholds, and generate high-quality information on CIC numbers and frequencies, clone sizes, linkages across sites and classify clones based on their extent of metastatic activity. Using of CIC-MM plot, statistical models and machine learning approaches, we generated high-resolution clonal maps of metastasis for each animal. The information generated included clone types and system-wide metastasis, similar and dissimilar clonal patterns of dominance at heterotopic sites and their routes of metastases. The data revealed previously unknown influences of cellular genotype and their implanted sites on selecting certain clones with specific system-wide clonal patterns, and identified rare LRRC15 expressor clones (classified as CIC.Toti) predisposed to exploit ‘all’ sites, albeit at varying degrees of dominance. The genomic technology and computational methodology described here are tissue-agnostic. They enable rapid adoption for an investigation into various stages of system-wide metastasis and growth of transplantable malignant cells at the highest clonal resolution.

## Introduction

Development of metastases is the leading cause of cancer fatality and preventing the development of metastases or treating such lesions once developed is a primary aim of systemic treatments (1, 2). Processes that result in the development of clones with metastatic potential within the primary tumors and the determinants of their colonization of heterotopic sites remain poorly understood, and likely differ significanlty among patients, limiting efforts to develop precision therapy (1). Thus, the capacity to quantitatively analyze metastasis at a clonal level across at different organ sites and traces the sources of inter-relatedness and growth potential of these lesions would provide insights that point to the development of precision therapies.

Cellular DNA barcoding is emerging as a powerful next-generation sequencing (NGS)-based genomic technology for clonal tracking *in vivo* and *in vitro* (3–9). Several ‘proof-of-concept’ studies are available in the literature to suggest that this technology can be used to understand normal stem cell (5, 10–12) and cancer cell heterogeneity (3, 4, 13–18), therapy resistance (7, 8, 13, 14, 19) at clonal resolution. However, the lack of standardized analysis, visualization and reporting of barcode data poses a significant challenge (5, 10, 12, 20, 21).

Herein, we present a novel NGS-based genomic barcoding solution to enable tracking large numbers of single cells *in vivo* to quantify the clonal growth and their metastatic spread. Specifically, we demonstrate the application of this technology using barcoded competitive clone-initiating cell (BC-CIC) assay of high-grade serous ovarian cancer (HGSOC) cell lines to understand the effects of LRRC15 expression and primary tumor sites on the metastatic spread. HGSOC patients typically present with metastatic disease and have a poor clinical outcome. Actionable targets mechanistically linked with system-wide metastatic progression of HGSOC have not been well described. A recent report demonstrated actionable type-1 transmembrane LRRC15 gene as significantly overrepresented in bowel metastasis compared to their matched primary ovarian tumors. This gene was suggested to play a role in omental metastasis (22). The clonal analysis of BC-CIC assay revealed previously unknown influences of LRRC15 expression and implant site on clone-initiating cell (CIC) frequencies, clone sizes and patterns of metastasis. The data also suggested that metastasis to local peritoneal sites likely uses a different route compared to metastasis to distal tissues. The classification of CICs based on their extent of system-wide metastasis has provided a complementary framework to understand clones that needs to be targeted for therapeutic benefits.

## Results

### Multifunctional high-complexity lentiviral DNA barcoding libraries

To generate technology that would simultaneously permit clonal tracking, *in vivo* imaging, library identification by fluorescence and sequence, and the ability to target of barcoded cells via antibody tags, we engineered multifunctional barcode library plasmid vector backbone (Figure S1A). Our lentiviral DNA barcode libraries have the following features a) MNDU3 promoter driven expression of red-shifted firefly luciferase (Luc; validation in Figure S1C) fused to a fluorescent protein (FP) gene (mRuby3, mTagBFP2, T-Sapphire, mWasabi or EGFP; the first 4 can be readily unmixed by FACS (Figure S1A; validation in Figure S1D) via a “self-cleaving” P2A linker peptide sequence. This MNDU3-Luc-P2A-FP cassette ensures equimolar expression of Luc and FP via a ribosomal skipping mechanism and allows both identification by flow cytometry via the FP as well as in vivo tracking of tumor spread via the Luc; b) PGK promoter driven expression of a human influenza hemagglutinin (HA) tagged sodium/iodide symporter (PGK-HA-NIS) gene thus allowing cells to be targeted by antibodies (validation in Figure S1E; not exploited in the present study). Next, we introduced a scaffold downstream of MNDU3-Luc-2A-FP cassette to clone non-coding semi-random barcodes as described elsewhere (10). In order to expand the repertoire of DNA barcode libraries, we added a unique 4 bp index sequence (index 1; Figure S1A) to the 5’ end of the barcode sequences and adapted the forward and reverse primer binding sequences to our design (adopted from (7)) for PCR based amplification of barcode sequence and NGS library preparation. The possible theoretical estimate of unique barcodes in our libraries is 4.19 x 10^6^ and the estimates based on transformed bacterial cell plating ranged between 1.1 x10^6^ – 3.0 x10^6^ (n=25 libraries, median diversity of 2.1 x10^6^) barcodes and the titer of concentrated lentiviral supernatants was in the range between 3.9 x 10^12^ - 2.8 x 10^13^ (median titer of 2.1 x 10^13^) transduction units/ml (Figure S1B). Barcode analyzed from plasmids isolated from 250 individual bacterial clones showed no redundancy (data not shown). In this present study, we assigned library 1.2 to OVCAR5-shLRRC15 (referred to as LRRC15-) and library 1.4 to OVCAR5-shControl (LRRC15+) cell lines (Figure 1A) and performed competitive tumorigenesis assays in mice (Figure 1B) using two different orthotopic xenotransplant methods (Figure 1C).

**Figure 1.**
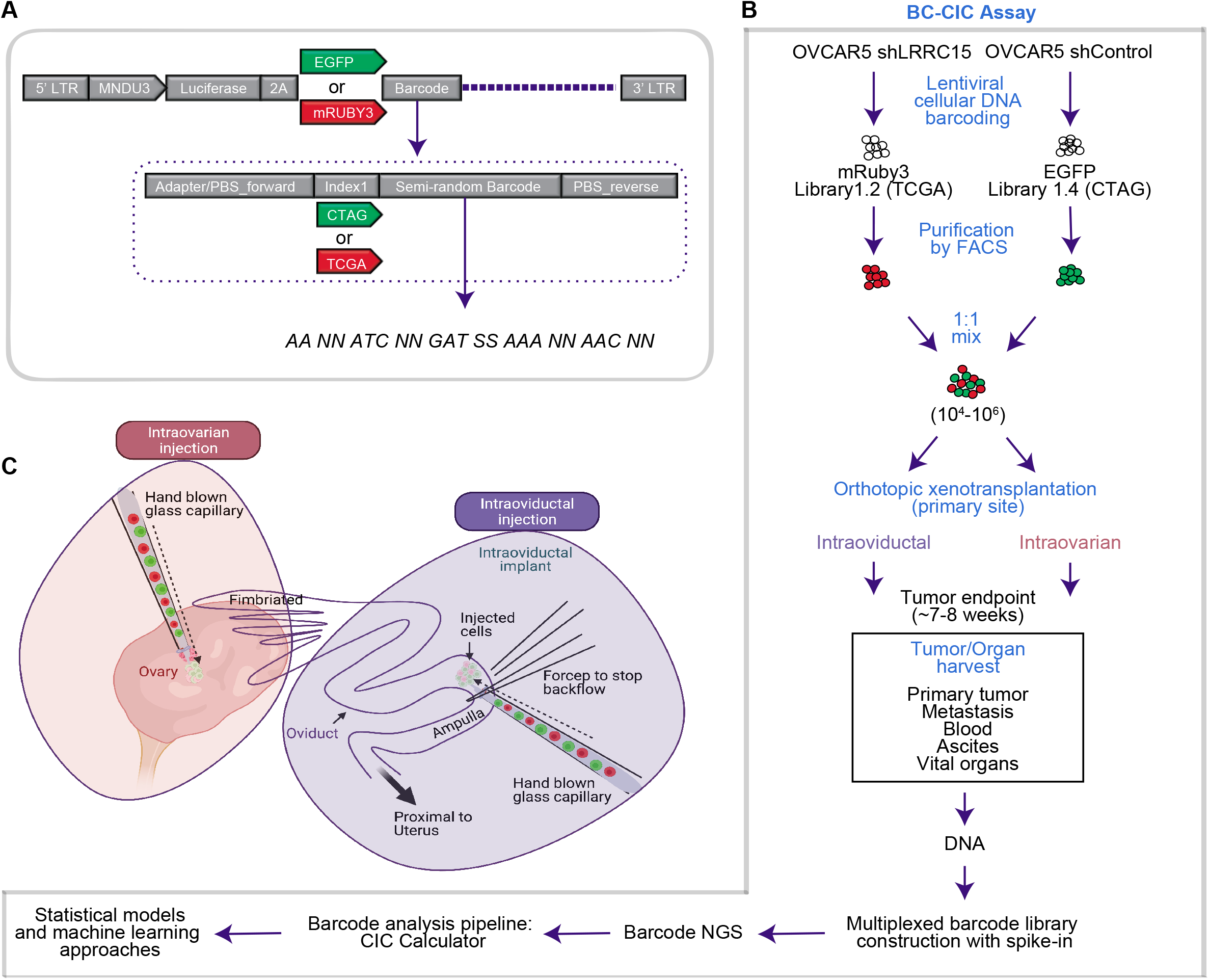
Barcoded Competitive Clone-Initiating Cell (BC-CIC) Assay. **A)** illustration showing EGFP_index 1.4 and Ruby3_index 1.2 barcode libraries used in the study. **B)** experimental design for BC-CIC assay. OVCAR5-shControl and OVCAR5-shLRRC15 were tagged with EGFP_index 1.4 and Ruby3_index 1.2 libraries and FACS sorted. The FACS sorted cells were mixed in equal numbers and co-transplanted at intraoviductal or intraovarian sites in NSG mice. Mice were monitored for tumor development till study endpoint. At necropsy primary tumor and tissues/organs were harvested and DNA was extracted. Multiplexed sequencing libraries were prepared with “spike-in” control barcodes by PCR and subjected to NGS. The data was analyzed using CIC Calculator. **C)** illustration of intraoviductal and intraovarian injection procedures in mouse.

### Spike-in control barcodes and multiplexed library preparation

To control variables associated with NGS sequencing library preparations and to set a threshold for data filtering steps, we additionally created a small library of 17 known barcodes (Sanger sequenced identified; Figure S2) plasmids of index 1.6 (Figure S1A) mixed at precise copy numbers (Figure S2A) to serve as internal calibration control. The “spike in” barcode control library was added to each sample during PCR based amplicon sequencing library preparation step (Figure S2B). Furthermore, the sequencing library preparation approach enabled labelling of each sample with 1 of the 20 unique indices (index 2.1 to index 2.20) and multiplexed per sequencing lane on Illumina HiSeq 2500 sequencer (Figure S2B), thus decreasing total sequencing costs.

### Implementation of Clone-Initiating Cell (CIC) Calculator

The flow chart of CIC Calculator is shown in Figure 2A. Briefly, indices were first identified from each read, unique barcodes from each library identified and counted, then barcodes from the same library with ≤2 mismatches pooled as likely sequencing errors (5, 11). This yielded count tables per unique index. We noticed that a large portion of the barcode clones were small clusters with less than 10 reads. Each sequenced flow cell contained ~500K unique barcodes. Low read number (<10 reads) clones took up to 90% of unique barcodes while accounting for only 0.5% of total read counts. Such small clones are assumed to be technical noise, i.e., contaminations introduced from PCR or NGS. Receiver-operator characteristics (ROC) based on the known ‘spike-in’ controls were used to filter out such technical artefacts (Figure 2B;). We observed a log-linear relationship between the known copy numbers in the spike-in control barcode library and reads with the exception of the single copy input, as one would expect due to Poisson sampling at limiting numbers (Figure 2C). As such, single-copy spike-ins were removed from the analysis which resulted in an improved R^2^, as well as better sensitivity (Figure 2D, Table S1).

**Figure 2.**
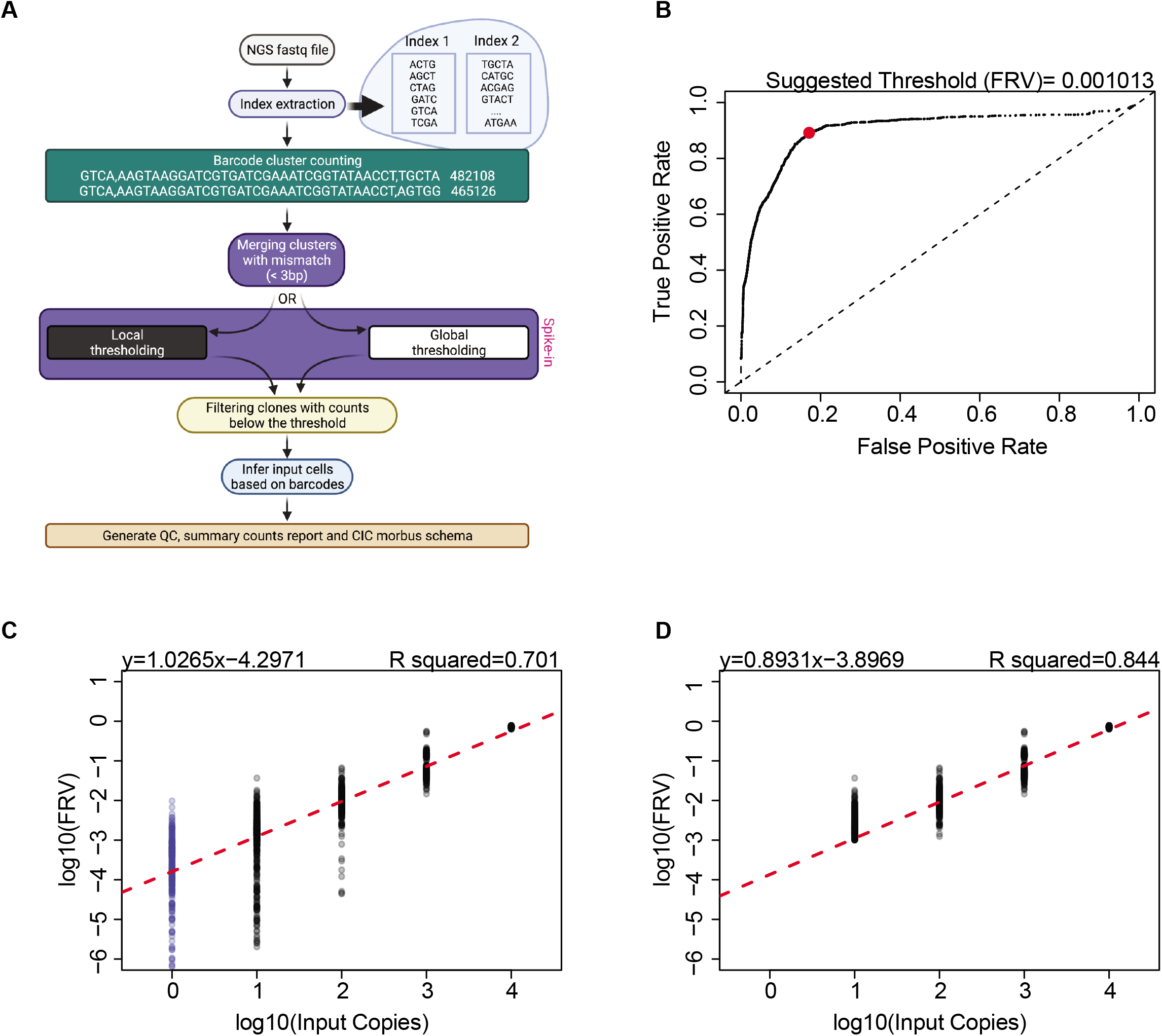
Clone-Initiating Cell (CIC) Calculator. **A)** flow chart showing the data deconvolution steps following NGS sequencing. **B)** receiver-operator characteristics (ROC) based on the known ‘spike-in’ controls were used to filter out artefacts. **C)** plot showing log-linear relationship between the known copy numbers (1 to 10,000) in the spike-in control barcodes and sequencing reads. **D)** removal of single-copy spike-ins improved the R^2^ values.

The CIC Calculator provides two different strategies for the clone size thresholding: local sample thresholding and the global sample thresholding (see flowchart, Figure 2A). By design, each sample (identified by a unique index 2) contains its own dedicated spike-in control barcodes. The local sample thresholding calculates the clone size threshold using only the spike-in in the specific sample of interest, which is more accurate and specific to the sample. The global sample thresholding instead pools all spike-in control barcodes across a group of samples from the same experiments, calculating a shared threshold for all samples. This is specifically useful if and when ROC curve might fail in some samples for various technical reasons. A stable ROC performance can be achieved for all samples in the same experiments. The two strategies can be set by the user before execution. The CIC Calculator was implemented with R and python. Codes will be available through https://gitihub.com.

### Growth characteristics of barcoded HGSOC cell lines in vitro

For the competitive tumorigenesis assay development, we exploited the OVCAR5-shControl and OVCAR5-shLRRC15 models (Figure 3A). LRRC15 was recently discovered as a top gene positively associated with bowel metastases in ovarian cancer patients and genetic or pharmacological targeting of this gene led to reduced metastasis of OVCAR5 cells (22). Herein we barcoded OVCAR5-shControl and OVCAR5-shLRRC15 cells, cultured for 48 hr for fluorescent reporter expression and FACS purified for xenotransplant studies discussed below (Figure S3A-B). In clonogenic assays, colonies obtained from OVCAR5-shLRRC15 were significantly smaller compared to OVCAR5-shControl (Figure 3B, Figure S3C), however, no significant difference was observed in clonogenic efficiency between the two cell lines (Figure 3C).

**Figure 3.**
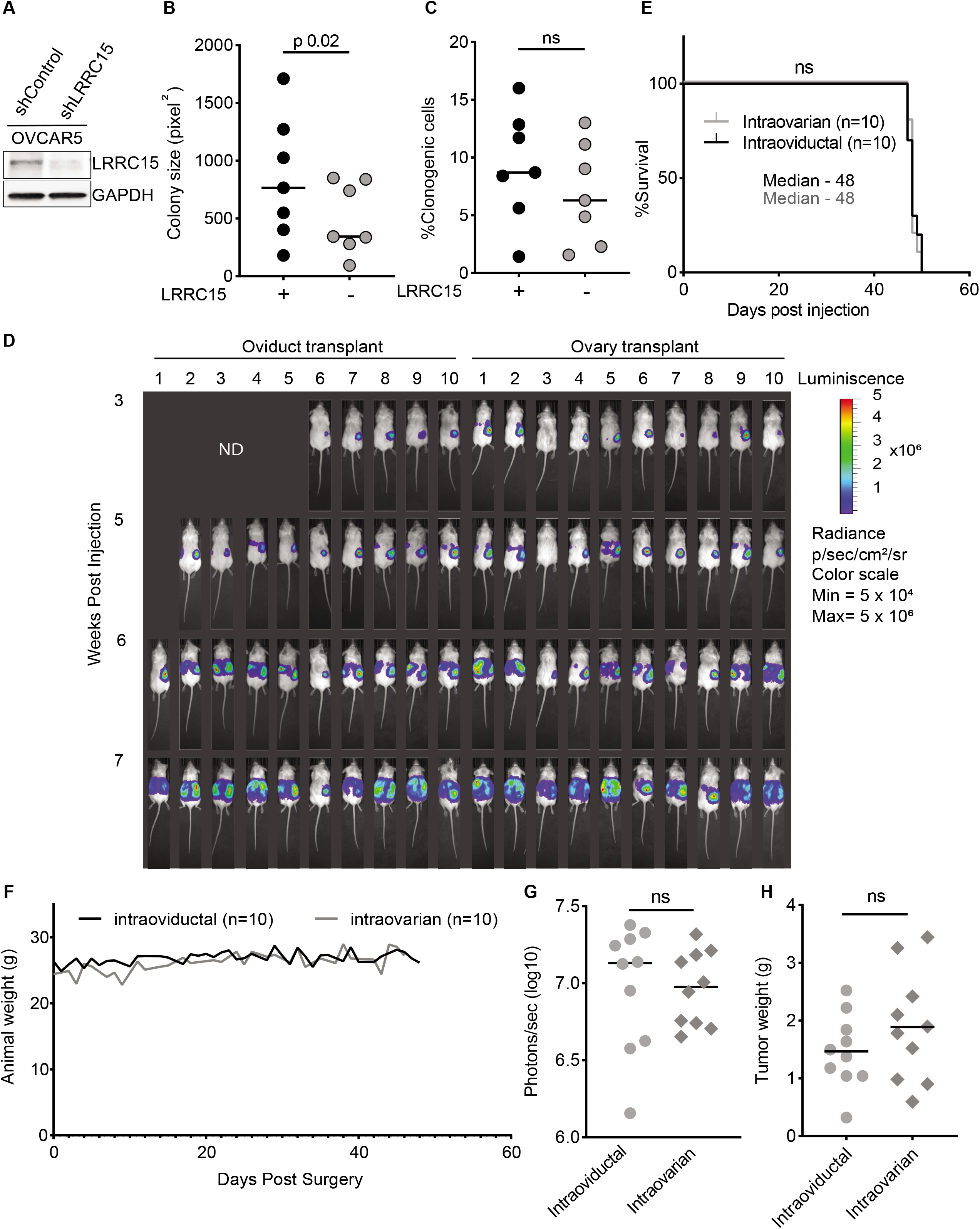
In vitro and in vivo growth characteristics of OVCAR5 cells. **A)** western blot analysis of LRRC15 levels in OVCAR5 derivatives. **B)** colony size (paired student t-test). **C)** clonogenic cell frequency. **D)** bioluminescence imaging of co-transplanted mice show the growth of tumors in NSG mice. **E)** Kaplan-Meyer analysis of overall survival. **F)** weight of NSG mice post-cell transplant. **G)** bioluminescence photons emitted in tumor bearing mice at ~7 weeks. **H)** correlation between dose of transplanted cells and tumor weight. Student t-test was performed to determine significance.

### Development of intraovarian and intraoviductal orthotopic xenotransplant models using highly immunodeficient NSG mice

To understand how the microenvironment influences the competitive growth of HGSOC clones we chose two different orthotopic sites, ovary and oviduct. These are prominent sites of ‘primary’ growth of tubo-ovarian cancers. We established robust surgical procedures in ~8 week of age highly immunodeficient NOD/SCID/IL2rγ-/- (NSG) mice to inject HGSOC cell lines under ovarian bursa (intraovarian site) or within ampullary ducts of oviduct (intraoviductal) using glass needles (see illustrations in Figure 1C, Figure S4A-B) with inner diameter of ~150 μm (23) (confirmed microscopically) (Figure S4A). All animals survived the surgical procedure irrespective of the site of injection or cell numbers injected and showed 100% tumor take for both intraoviductal (10/10) and intraovarian (10/10) injections as evidenced by time course bioluminescence imaging of transplanted animals (Figure 3D) and at necropsy (Figure S5).

### Characterization of tumors by conventional approaches shows no effect of injection site on transplanted ovarian cells

We monitored the co-xenotransplanted mice and sacrificed them once they showed distended abdomen, inability to ambulate, loss of body weight and/or inability to feed. All mice (two groups: intraoviductal and intraovarian) came down with aggressive disease and were euthanized around 7-8 weeks (median survival was 48 days for both groups). Between the two groups, no difference was noted in survival (Figure 3E) or animal weight (Figure3F). The groups also showed, similar bioluminescence signals (Figure 3G), tumor weight (Figure 3H), and organ weights (Figure S6A-H). Necropsy revealed tumors at injection sites (primary tumors and in omentum, bowel and other organs in both the groups (Figure S5). Transplanted total cell dose and weight of primary tumors were not correlated (Figure S6I). The conventional methods did not have the resolution to distinguish growth and metastasis of co-injected OVCAR5 cells in these injection sites. We designed primers targeting lentiviral vector to PCR detect transduced cells (see library construction in Methods for details). Gel electrophoresis analysis of PCR products of genomic DNA from primary tumors, omental metastatic tumors, ascites, blood and various organs showed system-wide metastasis in both groups (Figure S7).

### Barcoded Competitive Clone-Initiating Cell (BC-CIC) assay identifies major and consistent influence of genotype on CIC frequencies across sites

To investigate CICs of HGSOC derived cell lines, we performed BC-CIC assays (Figure 1B). Briefly, ‘barcoded’ OVCAR5-shControl (Index 1.2) and OVCAR5-shLRRC15 (Index 1.4) cells were mixed in equal proportions (10^4^-10^6^ cells; see Table S2) and co-transplanted into either right oviduct (n=10, mean = 355082 cells) or right ovarian parenchyma (n=10, mean = 437128 cells) in NSG mice per above. Genomic DNA was isolated from various anatomical sites including primary tumors and barcode sequencing libraries were constructed as described above. Amplicon libraries from barcode signal ‘positive’ sites, independently verified by gel analysis (Figure S7), were prepared with “spike in control barcodes”, multiplexed and sequenced (Figure 1C). Clones were identified and analyzed using CIC Calculator (Figure 1C). Given the distinct mechanisms of metastatic spread observed, we next investigated whether this relationship was underpinned by reproducible clonal patterns and whether there was any relationship between these, primary tumor site, or LRRC15 status. As a first pass, we looked at whether the total clone number per mouse was affected by either LRRC15 or injection site. Indeed, we observed a highly significant effect of LRRC15 knockdown on total clone numbers as well as a modest effect of injection site (Aligned Rank-Transform (ART) ANOVA p=2.1547×10-7 and 0.036 respectively (Figure 4A). Around 80% (range 71%-87%) of CICs in intraoviductal and ~90% (range 80-94%) of CICs in intraovarian group displayed metastatic behavior and greater heterogeneity in % of metastatic CICs induced by LRRC15 knockdown (~30% standard deviation vs ~5% in shControl CICs in both groups) (Figure 4B; Table S3).

**Figure 4.**
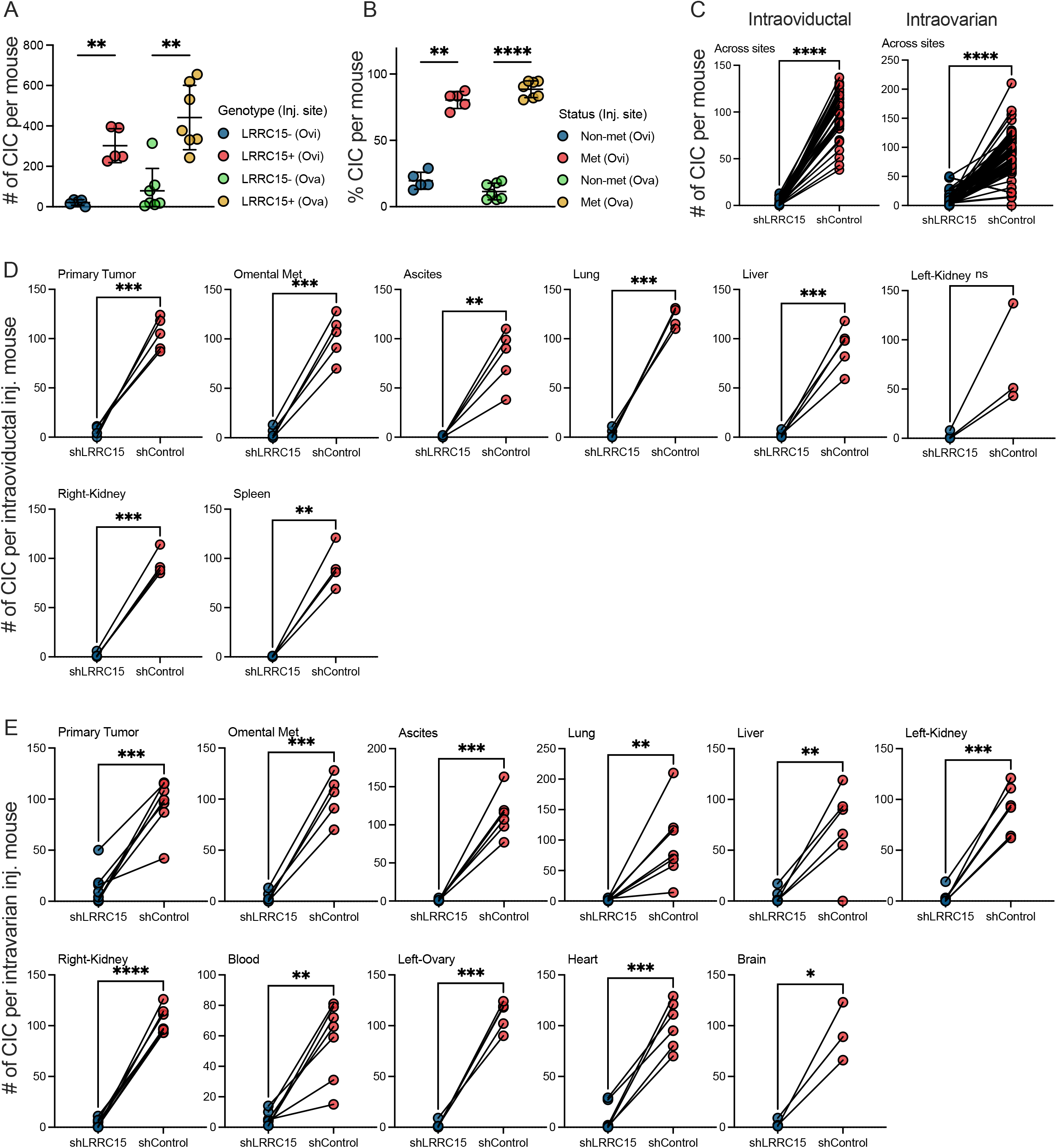
CIC numbers across sites are influenced by LRRC15 gene. **A)** total CIC per mouse split by LRRC15 status in both groups. **B)** total metastatic and non-metastatic CICs per mouse split in both groups. Groups in panel **A** and **B** were compared using one-way ANOVA and Tukey test, **<0.005 and ****<0.0001. **C)** CIC numbers across sites in both groups. **D)** CIC numbers in each site in intraoviductal co-injected group. **E)** CIC numbers in each site in intraovarian co-injected group. LRRC15- and LRRC15+ groups were compared using paired two-tailed t-test, *<0.05, **<0.005, ***<0.0005 and ****<0.0001.

We find irrespective of the injection site, LRRC15 knockdown dramatically reduced CIC frequency of OVCAR5 cells across all sites (Figure 4C). For example, in intraovarian co-injection studies, mean CIC number of OVCAR5 shControl cells at primary site was 95 and of shLRRC15 cells was 14 (mean difference 81; paired student t-test p<0.0005) (Figure 4D). We report similar trends for all organs for both groups (Figure 4D-E).

The maximal estimated clonal output from CICs across sites (corrected for total tissue/organ weights) in intraoviductal group (n=5 mice) identified the largest LRRC15-clone (44,886 cells) in primary tumor of mouse #48 and largest LRRC15+ clone (1,229,023 cells) in left-kidney in the same mouse. In intraovarian cohort, the largest LRRC15-clone (188,377 cells) was detected in cardiac blood of mouse #60 and largest LRRC15+ clone (2,077,136 cells) in omental metastases of mouse #62 (Table S4). Clones dominating across anatomical sites in both orthotopic xenotransplant models clearly demonstrated that the superior fitness of CICs was dictated by genotype (LRRC15 expressors over knockdown) (Figure S8 and Table S4).

### Lowest injected cell dose is saturating

We next analyzed the total number of barcodes detected per mouse, and the number of mice that expressed each barcode. Notably, the ~90% of barcodes in both libraries were detected only in a single mouse (Figure 5A), as expected, given that each library includes over one million unique barcodes. In each library, however, ~10% of barcodes were detected more than once (Figure 5A). Based on the library size and number of clones observed from each library, such multiple detection is unlikely to have occurred by Poisson sampling (p<<0.001), and thus these barcodes must have been over-represented in the libraries (with probable numbers ranging in the ~100s-1000s of copies in library 1.4, and ~10000-50000 copies in library 1.2). However, we did not find any significant relationship between injected cell numbers and the number of observed barcodes (Figure 5B). This lack of correlation suggests that tumors likely saturate at injected cell numbers well below even the lowest tested. Whether this is due to the technical limitation of sampling, or that the mice saturate at a lower number of tumor initiating cells, however, still needs to be determined. This is, however, an important consideration in experimental design as most studies inject cell numbers at or in excess of those tested here (22), and thus may be over-loading the biological assays system.

**Figure 5.**
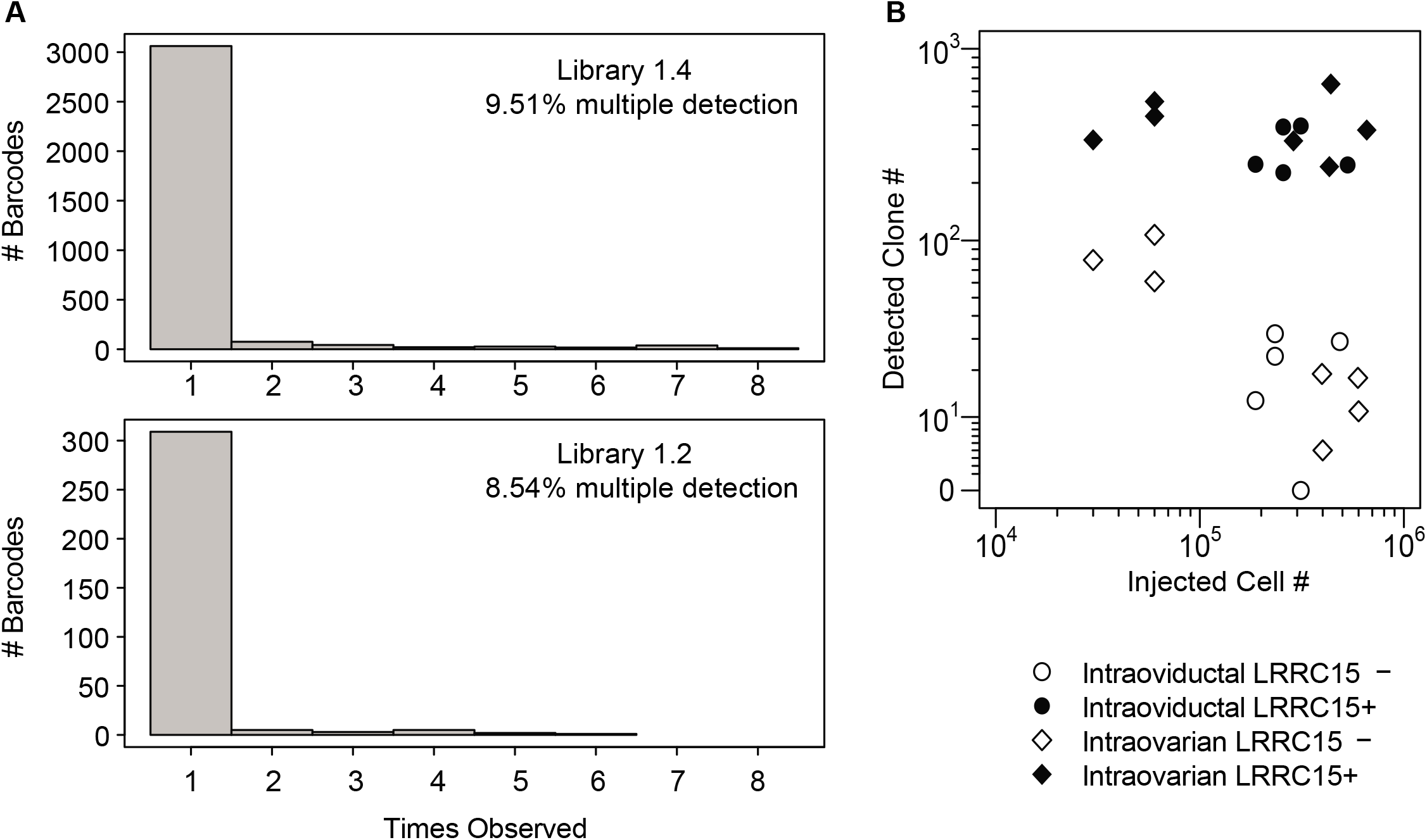
Barcode libraries are reasonably diverse, but input cell numbers are an important consideration in experimental design. **A)** number of mice in which each barcode was detected is shown for the 1.4 (top) and 1.2 (bottom) libraries. The % of barcodes (of total clones) which appeared in more than one mouse is indicated in the top right for each library. **B)** the number of clones detected per mouse is shown compared to the cell number injected. Injection site is indicated by point shape and LRRC15 status (“+” = wildtype, “-” = shRNA knockdown) is indicated by point fill as shown in the legend (right). Axes are asinh(x/10) transformed (equivalent to a biexponential transform) as relationships are expected to be logarithmic but with 0 values present.

### CIC Morbus-Mandala (MM) plot identifies influence of genotype on complex metastatic spread across sites at clonal resolution

CIC-Calculator utilizes a novel chord diagram named CIC Morbus-Mandala Plot that provides a visual description of clonal evolution of cancers, including inter-relationships among metastatic deposits. The external circular ring presents the tumors harvested in various organs (color table shown in Table S5). The lines in the sectors of the internal ring represent the presented barcode clones in the corresponding anatomical sites. A link will be drawn between two sectors if the same barcode is found presented in both sites. The circular layout makes it germane to visually comprehend complex relationships between metastatic CICs and various tissue/organ sites.

There are various types of CIC-MM plots described here and they are helpful to depict the following information pictorially. 1) CICs from multiple or individual experiments conducted in a ‘single’ mouse (see examples in Figure 6A-B); 2) depiction of system-wide linkages per clone (Figure 6A-B); 3) video depiction of system-wide clonal spread in an organism (Video S1). In order to understand the clonal metastatic behaviors represented in CIC-MM plot, we categorized CICs into four types namely a) CIC of indeterminate potential (CIC.IP, present only in primary site), b) CIC of mono-metastatic potential (CIC.Mono, observed only in one site +/- primary), c) CIC of pluri-metastatic potential (CIC.Pluri, observed in more than one but not in all sites), and d) CIC of toti-metastatic potential (CIC.Toti, present in ‘all’ sites). Examples of clones representing each of these categories are presented in Figure 7A.

**Figure 6.**
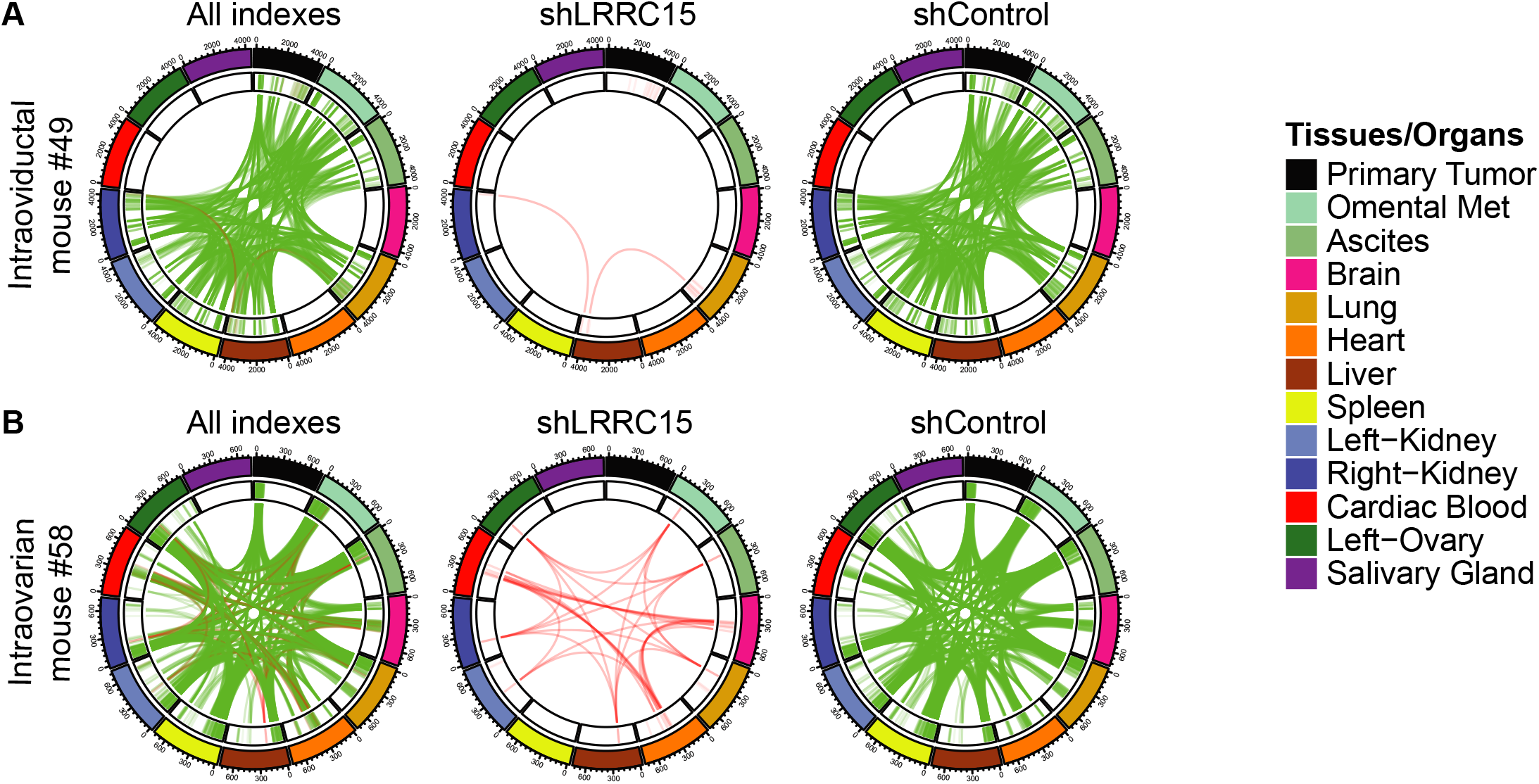
CIC-Morbus Mandala (CIC-MM) plots pictorially depict system-wide metastasis. **A)** a representative CIC-MM plot depicting system-wide metastasis of CICs in mouse#49 co-injected with barcoded OVCAR5-shControl and OVCAR5-shLRRC15 cells in intraoviductal site. **B)** a similar representative CIC-MM plots of CICs in mouse #58 co-injected in intraovarian site. Plot depicting influence of LRRC15 status on CIC.Pluri class.

**Figure 7.**
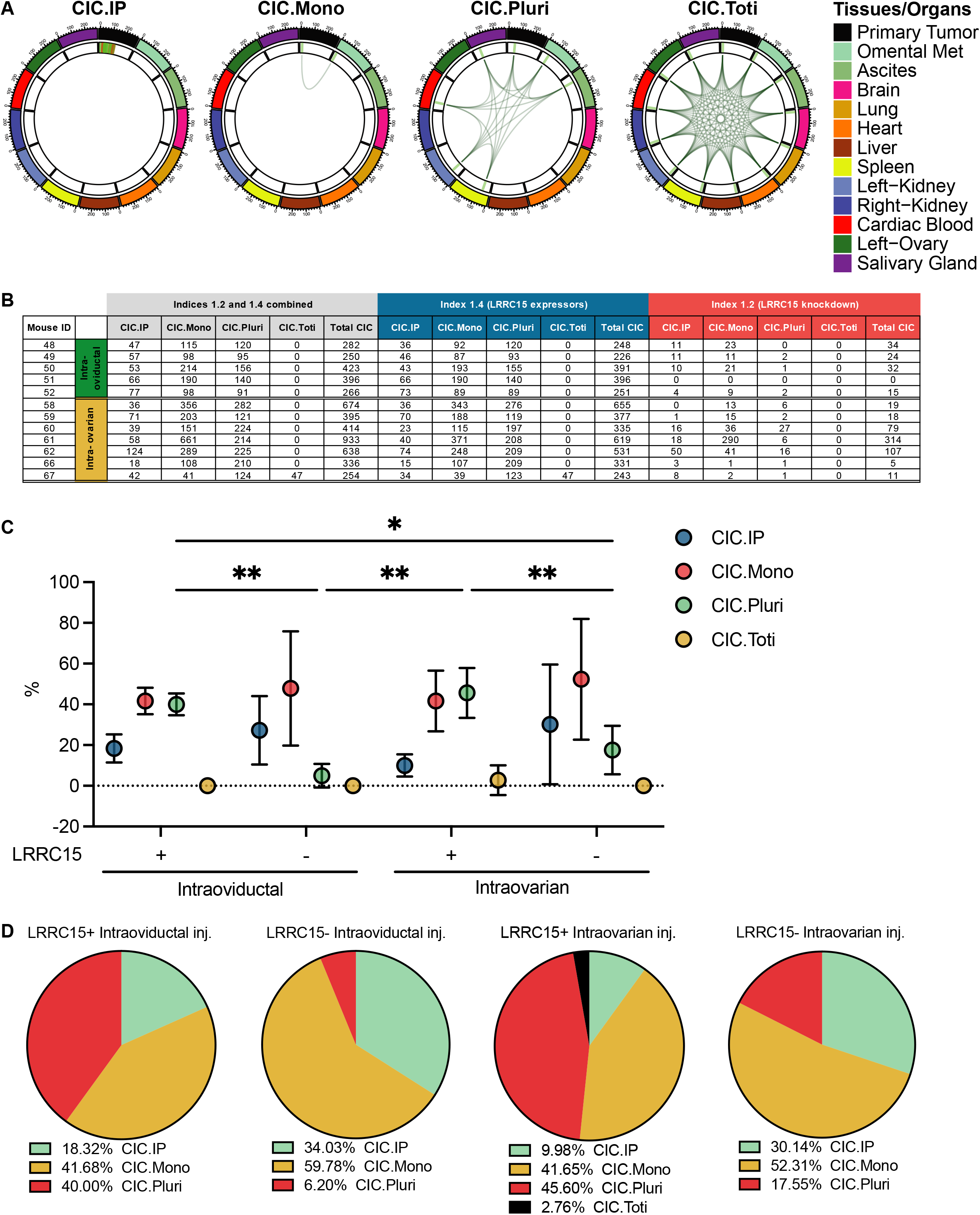
CIC classes and genetic influence. **A)** representative CIC-MM plots of CIC.IP, CIC.Mono, CIC.Pluri and CIC.Toti. **B)** CIC numbers per class split by LRRC15 status and injection site. Groups compared using two-way ANOVA and Tukey test, *<0.05, **<0.005 and ****<0.0001. **C)** venn diagram showing % of CIC classes. Note: CIC.Toti class was observed only in LRRC15+ CICs in a single intraovarian injected mouse.

The data from BC-CIC assays (co-injection of LRRC15 expressors vs silenced) in 12 mice (5 intraoviductal vs 7 intraovarian) using CIC-MM plot depicted the following. 1) The metastasis and growth of CICs in both intraovarian and intraoviductal sites led to similar complex system-wide diseases with notable inter-mouse differences (Figure S9 & S10). 2) CICs from both co-injected cell lines were represented system-wide. However, most CICs with organ/site linkages were from LRRC15 expressors clearly suggesting that targeting this gene largely neutralized OVCAR5 cell’s ability to metastasize and grow across sites. CICs are further resolved as percentage of CIC.IP, CIC.Mono, CIC.Pluri, and CIC.Toti (Figure 7B-D; Table S3).

### Metastasis to sites in the peritoneal cavity and distal sites is correlated with different routes

We next investigated whether there were any reproducible relationships between metastatic sites across mice. Interestingly, the relative relationships between metastatic sites appeared to be reproducible across mice and injection sites (Congruence Among Distance Matrices test p=0.001). When visualized in aggregate, these data revealed several distinct groupings of metastatic sites (Figure 8A,B). This included one group locally in the peritoneal cavity (local kidney, contralateral ovary, and omental metastasis along with ascites), another with distal sites (salivary gland, brain, heart, blood), and others which were intermediate between the two (lung, spleen, liver, contralateral kidney) (Figure 8A,B). Interestingly, the primary tumor was relatively distinct from all metastatic sites, suggesting that clone size in the primary site is not a good predictor of metastatic spread. These groupings were consistent between injection sites (Figure 8A-B). Importantly the observed relationships were highly consistent even when analyzed by individual mice as indicated by the CADM test (Figure 8B, Figure S11C-D).

**Figure 8.**
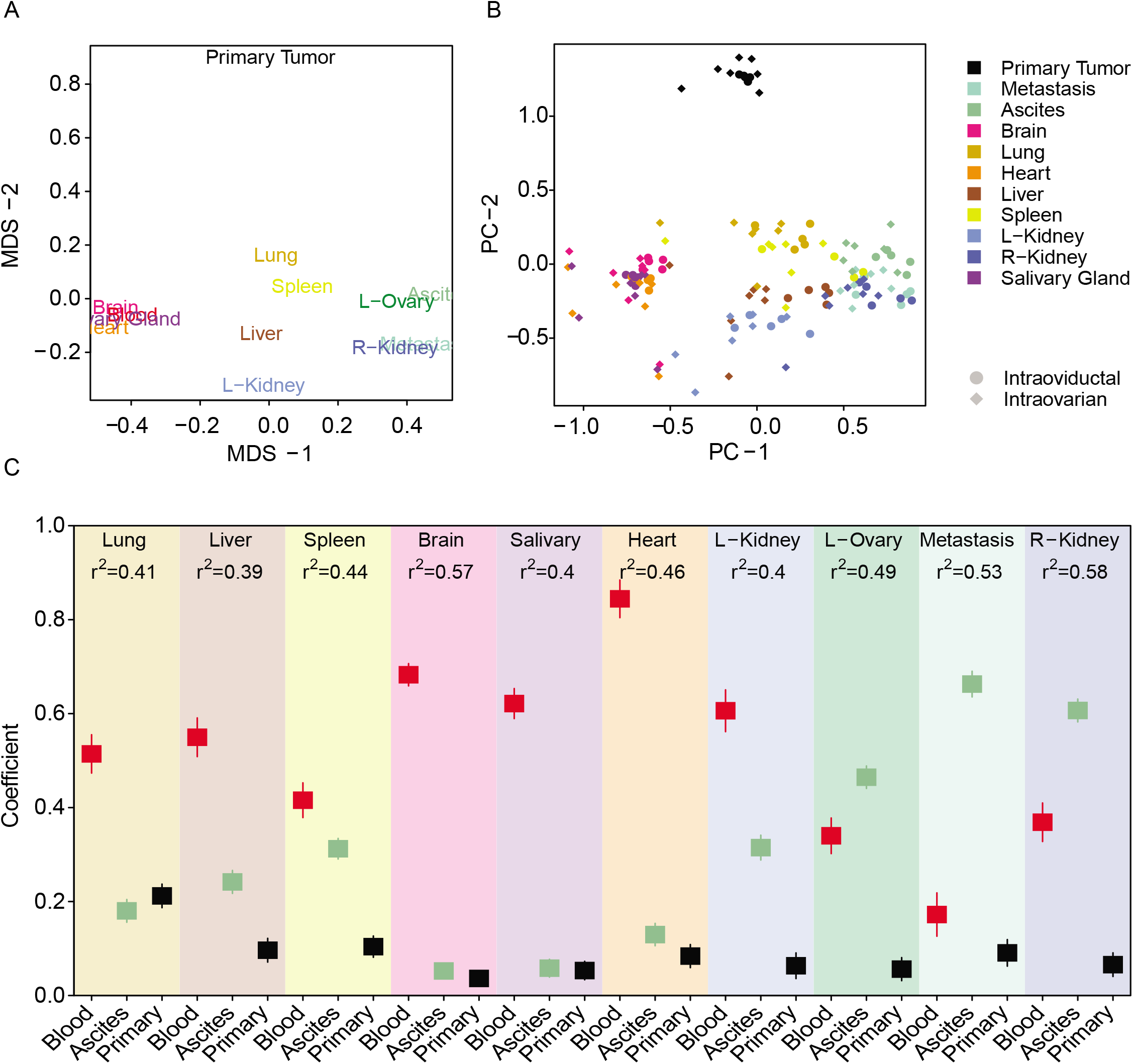
Clones detected in blood are associated with metastasis to different organs than those detected in ascites. **A)** relative relationships between sites of tumor growth. Multi-dimensional scaling (MDS) dimensionality reduction based on the mean distance matrix across mice is shown. Site names displayed at their embedded locations. The closer together sites are displayed on the plot, the more correlated their growth. **B)** per-mouse relationships between sites of tumor growth. The plot shows a PCA embedding based on the averaged distance matrix across mice (approximately equivalent to MDS) followed by embedding the per-mouse relationships into this space as a measure of how consistent these relationships are across individual animals. As with the MDS plots in (A), the closer the points, the more similar is their patterns of clonal growth. Injection site is indicated by point shape and site of growth is indicated by color as shown in the legend. Only sites measured in all mice are shown in this plot. **C**) coefficients of are shown for a series of multivariate linear regression models using clone size detected in the blood, ascites, and primary tumor to predict levels in each other site (all asinh(x/10) transformed to maintain linearity). Higher coefficients (values on the y-axis) represent a greater predictive power of that site for the overall clone size at the indicated site. Lines showing 2x the standard error are shown for each coefficient. See also, supplementary figures (S11-S12).

Since clone levels in blood were correlated with metastasis to distal sites, and clone levels in the ascites with peritoneal spread, we hypothesized that these might represent distinct mechanisms of metastasis. To quantify the relationship between blood and ascites to each metastatic site, we next constructed a series of linear models using clone abundance in these two sites along with primary tumor as an outgroup to predict spread to each other metastatic site. As expected from the earlier visualizations, distal sites were strongly and almost exclusively predicted by blood, while the local peritoneal cavity appeared to be predicted mainly through levels in the ascitic fluid (Figure 8C, Table S6, Figure S11E & Figure S12), with models explaining around half of the observed variability (r^2^ in the 0.4-0.6 range). Given that the current observations represent a snapshot at endpoint and thus early transit through these routes would be missed, these results demonstrate that clone abundance in blood and ascites can substantially predict metastatic spread.

### Multiple reproducible clonal patterns of engraftment exist and are influenced by injection site and LRRC15 status

We next performed a clustering analysis to identify clonal patterns which were reproducible across mice. Visualization of the identified clusters by various dimensionality reduction algorithms confirmed that the identified clusters appear to be relatively distinct and compact (Figure 9A). Importantly, all retained clusters were observed in 2 or more mice, with many being observed in most or even all mice suggesting that these represent common patterns of metastasis for the OVCAR5 cell line. These patterns included a very common cluster present in the primary site, but low or absent in all metastatic sites (Cluster I), another with prevalent peritoneal spread but no distal tissue contribution (Cluster IV), and another with distal spread and blood detection but relatively low peritoneal contributions which were observed in almost all mice with an intraovarian primary tumor, but none with an intraoviductal (Cluster VIII) (Figure 9B). A variety of other patterns were also observed including multiple in which large clones only at a single site (Figure S12). These likely represent an early emigration from the primary site. Interestingly, there was also a cluster with high levels at the blood-associated sites but without blood detection (Cluster X, Figure 9B), which may represent a clone that spread through blood at an earlier timepoint. Finally, approximately half of all observed clones were very small with no sites showing 10 or more cells (Figure 9B).

**Figure 9.**
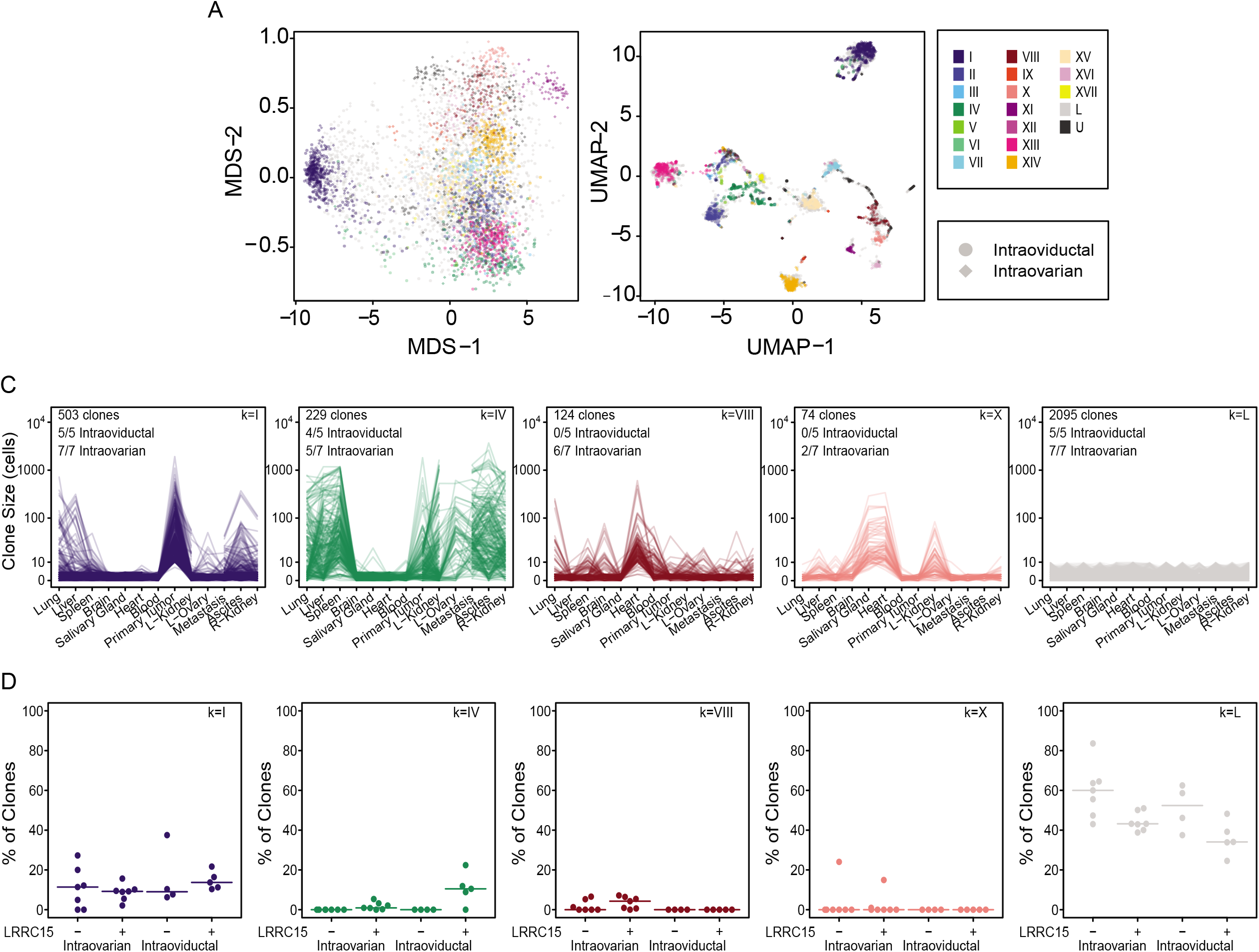
Common patterns of clonal growth exist and are influenced by injection site and LRRC15 status. **A)** the overall number of detectable clones is shown for each mouse separated by injection site and LRRC15 status (“-” = LRRC15 shRNA, “+” = LRRC15 wild-type). **B)** dimensionality reduction by MDS (left) and UMAP (middle) based on the pair-wise distance transformed Pearson’s correlations between all clones. Injection sites are indicated by point shape, and cluster membership is indicated by point color as indicated in the legend (right). **C)** clonal engraftment patterns are shown for selected clusters. Lines connect cell numbers detected for each clone between sites. Missing values for a given clone are shown as gaps between adjacent line segments. Clone numbers are asinh(cell #/10) transformed with the linear numbers indicated. The total number of clones which were part of a cluster, and the number of mice with at least one clone that was a member of that cluster for each injection site are shown in the top left. The cluster identifier for each is shown in the top right. **D)** the number of detectable clones which are part of selected clusters is shown for each mouse separated by injection site and LRRC15 status (“-” = LRRC15 shRNA, “+” = LRRC15 wild-type). See Figure S13 and S14 for engraftment patterns and injection site/LRRC15 comparisons for remaining clusters.

We next tested whether any of the observed clonal patterns were influenced by the primary tumor injection site or LRRC15 status. Interestingly, there was a significant enrichment of LRRC15 knockdown cells in the generally small clones (Figure 9C, ART ANOVA false discovery rate (FDR)=0.01). LRRC15 knockdown thus decreases both the total clone number, as well as the size of detected clones, suggesting a strong fitness deficit. Among most other clusters, however, clone frequency was roughly similar with a few notable exceptions (Figure 9C, Figure S13, Table S7). Interestingly, there was a significant enrichment for LRRC15+ clones with intraoviductal injection sites Cluster IV with spread throughout the peritoneal cavity (ART ANOVA FDR 0.01 for injection site, 0.004 for LRRC15 status, and 0.004 for the interaction of the two, Figure 9B-C). Finally, Cluster VIII with detection in blood and distal sites was significantly enriched in mice with intraovarian primary tumor site (ART ANOVA FDR= 0.02, Figure 9B-C, Table S7). Overall, these results demonstrate that LRRC15 knockdown gives a significant fitness deficit, though it doesn’t necessarily alter the metastatic potential of cells bearing it, and that primary tumor site may influence its available routes of metastatic spread and thus probable metastatic sites.

### Among the analyzed mice, ovarian tumors show higher competence for blood-based metastasis to distal sites

Given that blood and ascites appear to represent independent routes of metastasis, and within the mice analyzed we observed an enrichment of distal tissue metastasis from primary ovarian tumors. We next wanted to compare blood and ascites detection by primary tumor site. In the analyzed mice, however, we only had clonal measurements of blood in one of the intraoviductal mice. To circumvent this, we train an elastic net (EN)-based machine learning model to predict on a per clone basis whether or not it would be detectable in the blood. Following training on a subset of the clones from intraovarian mice, this model showed an overall per mouse median prediction accuracy of 0.92, the sensitivity of 0.71 sensitivity and specificity of 0.95 in the remaining clones from these mice on which it hadn’t been trained (ie. the validation set, Figure 10A and Figure S14A-B). Thus, predicted blood clones are very likely to be detected in the blood, though some clones which are detectable in the blood may be missed. One of the benefits of an EN model is that the final parameters contributing to each decision are explicit, unlike for more powerful but less interpretable deep learning approaches. This revealed that the primary indicators of a clone being detectable in the blood were a presence in the brain and heart (Figure 10B), two of the distal sites associated with blood from both in bulk (Figure 8), and in Cluster VIII (Figure 9C). Finally, when applied across all mice, we observed a significantly lower frequency of blood detectable clones in the intraoviductal compared to intraovarian mice (Wilcoxon rank-sum test p= 0.01, Figure 10C). This finding is further supported by the measured clone numbers in the intraovarian compared to the 0 observed and 0 predicted in the one intraoviductal mouse with measurements performed on blood (Figure S14B). Thus, among the observed mice, intraovarian primary tumors showed an improved competence for blood mediated metastasis.

**Figure 10.**
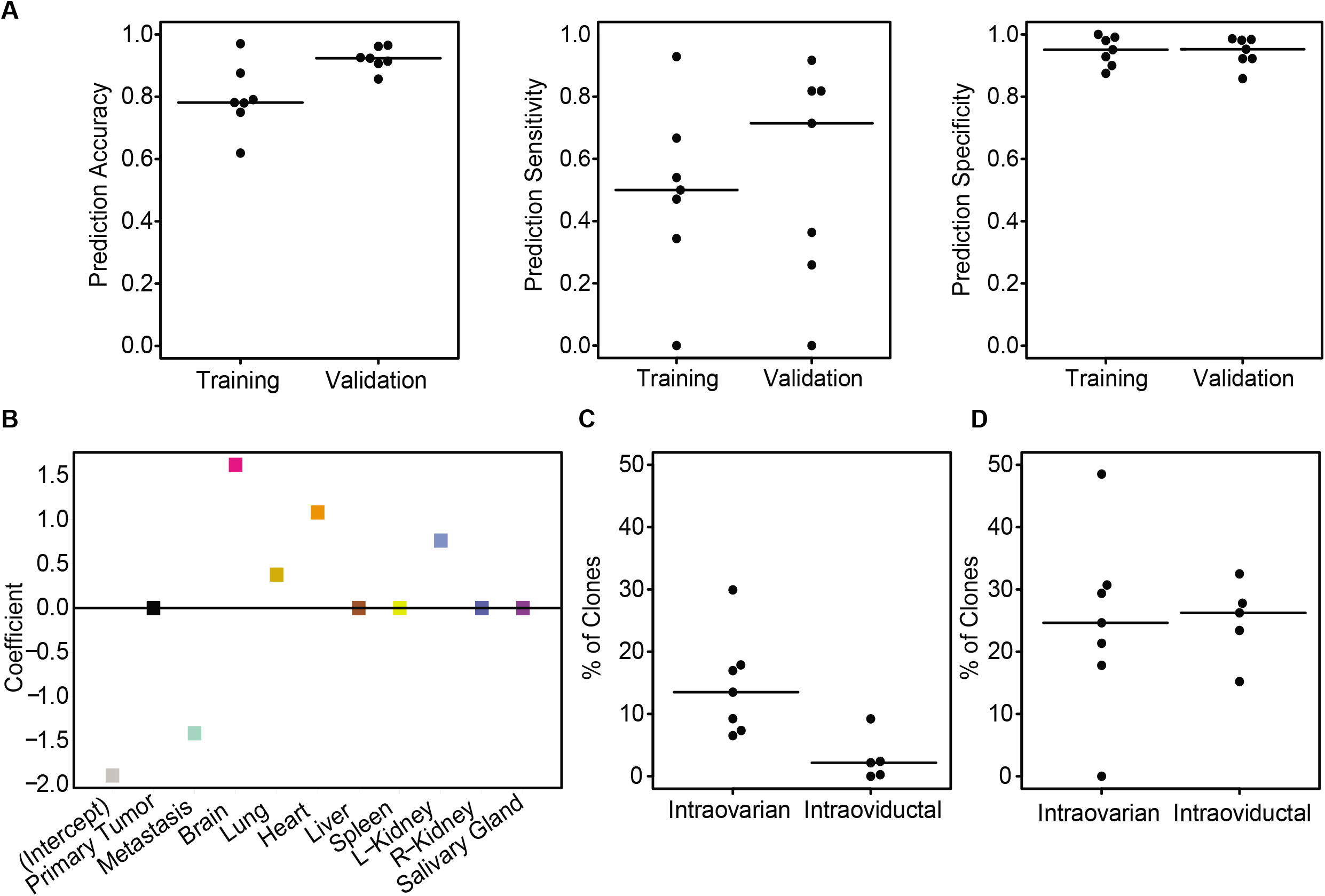
Primary tumors in the oviduct are deficient in blood metastasis those that arise in the ovary. **A)** overall accuracy (left), sensitivity (middle) and specificity (right) of an elastic net (EN) logistic regression to predict whether a given clone is present (detectable) in blood. The EN model was trained on 70% of blood detectable clones from each mouse in the intraovarian cohort and double that number of non-blood clones with remaining clones kept back for the validation set. Each point represents the predictions for clones from one mouse. Each plot shows the relevant metric for the training and validation sets. **B)** the coefficients retained for each site in the final EN model for predicting whether a clone will be detectable in blood or not and their magnitude. Positive numbers indicate a value that is used to predict that a clone is present in blood, while negative numbers indicate sites in which the presence there means the clone is less likely to be in blood. The negative intercept value represents the threshold of positives needed to indicate a clone is likely present in blood. **C)** percentage of clones per mouse predicted to be detectable in blood by the EN model separated by injection site. **D)** percentage of clones per mouse with measured detection in ascites separated by injection site. See Figure S14 for model hyperparameter fitting, measured percentage of clones detectable in blood for those mice which had measurements, and an EN predictive model for detection in ascites.

We also trained an EN model to predict whether a clone would be detectable in ascites (Figure S14C). However, this model proved less accurate with median validation set accuracy 0.82, the sensitivity of 0.40, and specificity of 0.86 (Figure S14D). This decreased accuracy could result from ascites mediated spread taking place in many cases earlier on than would be detected by our end-point measure. Predictions would also be further impacted by the fact that blood can mediate spread to many of these tissues and further increase the prediction difficulty. Despite the relatively lower performance, the model did still provide some useful information. Omental metastasis and the local kidney represented the primary predictors of detection in ascites based on this model (Figure S14E). While no significant differences were observed in ascites by injection site at the time of measurement (Wilcoxon rank sum test p=1, Figure 9D), the predicted ascites clones were significantly higher in intraoviductal tumors (Wilcoxon rank sum test p=0.01, Figure S14F). This observation fits with the observed enrichment in intraoviductal tumors for Cluster IV (peritoneal spread, Figure 9C-D). As size at metastatic sites reflects the entire history of metastasis, while the measurement is a snapshot at mouse endpoint, this may reflect a higher rate of metastatic spread via ascites earlier on in tumor progression.

Perhaps the most interesting results from the EN models come from a comparison of their coefficients. Omental metastasis was a counter predictor of blood detection (Figure 10B), while brain and heart counter predicted ascites spread (Figure S14E). As these were in both cases the primary positive predictors for the other type (brain and heart predicted blood, gut predicted ascites), this data strongly suggests that ascites and blood likely represent independent routes of metastasis.

### CIC-MM plot illustrates previously unknown genotype specific patterns of system-wide association of bowel macrometastasis (bowel met) clones

Finally, we analyzed the clonal composition in individual bowel met. At necropsy, mouse #61 transplanted with 120,000 cells at intraovarian site showed multiple overt bowel met. Such bowel met cause the terminal stage of HGSOC due to malignant bowel obstruction (24). We harvested 8 of these bowel mets and performed barcode analysis (Figure 11Ai). Clonal linkages using CIC-MM plot and clone size distribution provided fascinating and novel insight into the nature of system-wide metastasis and growth of bowel met clones (Figure 11A i-v). CIC-MM plots distinguished classes of LRRC15+ CICs and LRRC15-CICs dominating the bowel metastasis. In 8 bowel sites analyzed, CIC.Pluri class were observed for LRRC15+ in 8/8 sites and in contrast competing LRRC15-cells were CIC.Mono in 6/8 and CIC.Pluri in 1/8 bowel sites. The metastatic patterns of CIC.Pluri clones, irrespective of the genotypes, were nearly identical (Figure 11A i-v). The barcoded cells were ~10% of tumor cells in the bowel mets (Figure 11B). Analysis of clone size distribution indicated that both LRRC15+ and LRRC15-clones were equally competing in most of the bowel sites (Figure 11C-D). These unexpected findings provide an insight into behaviors of clones that potentially end life by compromising bowel integrity. The pattern of bowel metastasis is akin to multicellular attachment and growth but how LRRC15-cells attach to the bowel without cells of these clones undetected in ascites remains unclear. Overall, the findings point towards a model where lack of LRRC15 expression may reduce fitness of OVCAR5 cells to compete across primary and heterotopic sites except bowel wall.

**Figure 11.**
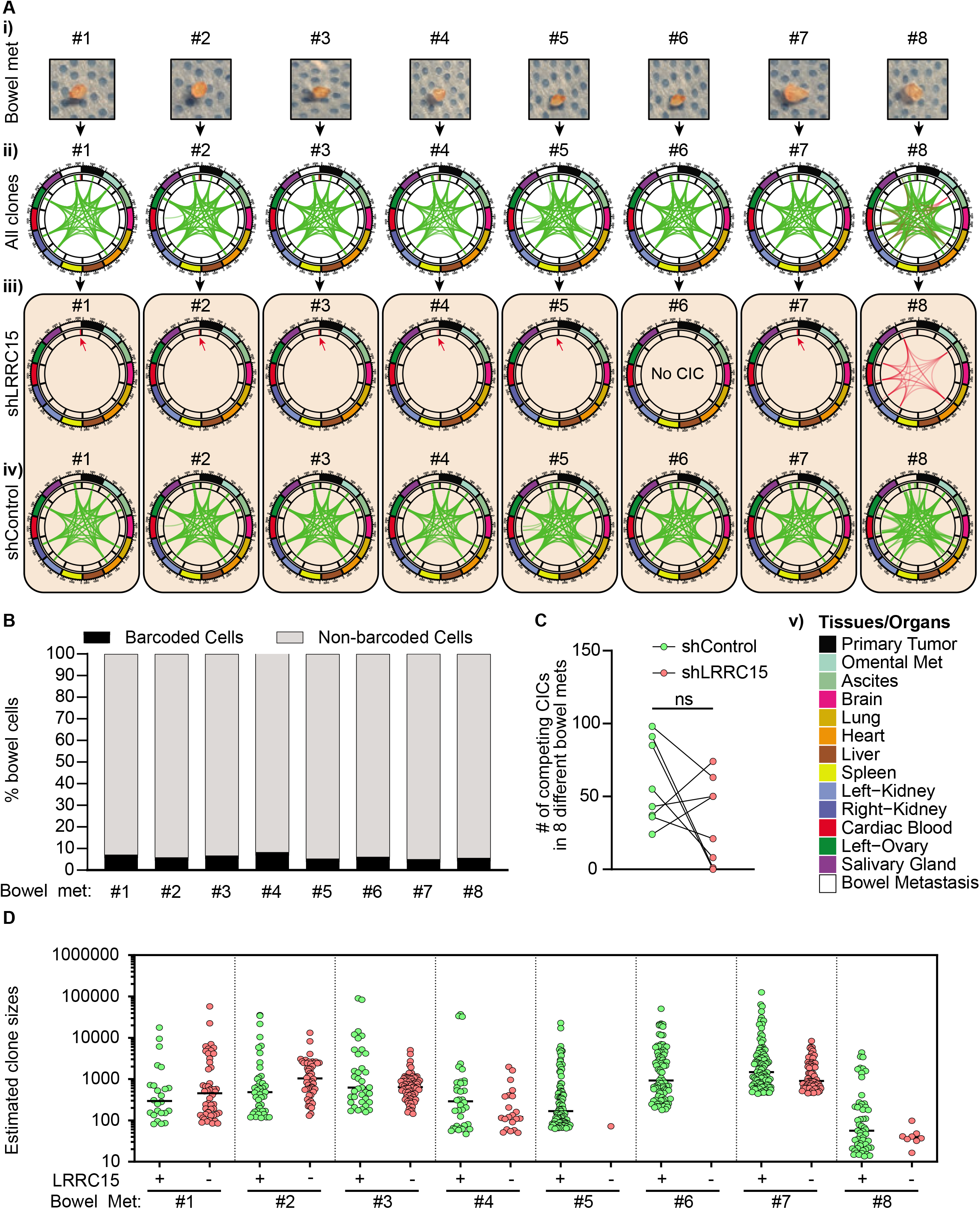
Clonal analysis of bowel macrometastasis (bowel met) and their system-wide linkages. **A i)** pictures of harvested bowel mets from mouse #61. **A-ii-iv)** MM plots depicting all clones, shLRRC15 and shControl respectively. **A-v)** color codes for tissues and organs. **B)** estimated percentage of barcoded cells in the bowel mets based on number of barcoded cells per μg of DNA. **C)** plot showing number of shControl and shLRRC15 CICs in bowel mets. D) Estimated clone sizes across in LRRC15+ and LRRC15-bowel mets.

## Discussion

We report development of cellular DNA barcoding based clonal tracking technology and end-to-end computational approach to infer system-wide metastatic cell clonal behavior and their complex relationships between primary and various heterotopic sites they metastasize and grow. Using BC-CIC assay design, we have demonstrated the power of cellular DNA barcoding technology and associated computational pipeline to analyze and visualize metastatic cell clonal data at highest resolution. The salient features of barcoding described here includes the following: 1) robust detection of clones (cells with shared barcodes). 2) quantification of number of clones in primary tumor and system-wide tissue/organ sites. 3) estimation of frequency of clones based on input cell numbers. 4) estimation of clone size based on internal spike-in pool of pre-selected barcodes of known counts and a choice to use local or global thresholding to determine clone detection limit. 5) simplified illustration of system-wide complex spatial linkages of clones using CIC-MM plots and finally, two distinct classification of system-wide spreading clones into behaviorally distinct and biologically meaningful types. The first approach involves classifying CICs into one of the four distinct classes: CIC.IP, CIC.Mono, CIC.Pluri, and CIC.Toti. The second approach uses a combination of dimensionality reduction and clustering to further dissect the biological meaning of the observed clonal patterns. We demonstrate that the CIC Calculator output data can be integrated with a suite of complementary statistical and machine learning approaches that uses CIC number, frequency, and system-wide clone size distribution as inputs from CIC Calculator to determine clone classes, their system-wide linkages and metastasis routes at clonal resolution. The demonstration of this technology/methodology is to standardize its application in cancer metastasis research.

Our methodology describes 1000s of clones and their growth patterns. Discussing such large numbers of clonal patterns can have practical limitations. The general clone classes (CIC.IP, CIC.Mono, CIC.Pluri, CIC.Toti), which are expressed as proportion of total clones, are biologically meaningful. For example, BC-CIC assay demonstrated that LRRC15 knockdown reduced CIC.Pluri and CIC.Toti suggesting the role of LRRC15 in influencing CICs with aggressive CICs. Standardized reporting of these CIC categories will enable comparison of findings from barcode studies across laboratories. Further insights into clonal mechanisms can then be made by subdividing clones into meaningful clusters in a biological system-informed manner.

During the early 1980s two important insights into metastasis processes emerged. The evidence was presented to claim the clonal nature of spontaneous melanoma cell metastasis by tracking X-radiation induced chromosomal rearrangements (25) and that melanoma cell metastasis may involve the natural selection of cells with preexisting properties to metastasize and grow at particular heterotopic site (26). Our statistical methods to study 1000s of clones and their growth patterns in heterotopic sites in various mice with input from the same pool of cells enabled us to look for such clonal patterns of preferred heterotopic sites. CIC clusters describe 17 different clonal patterns of system-wide metastasis. Given the parental OVCAR5 cell line was originally derived from patient ascites from highly heterogeneous high-grade serous ovarian cancer (HGSC) disease, such diverse growth patterns are therefore not surprising. Very interestingly, however, our results suggest that there are two distinct routes of metastasis, one through the ascites to the local peritoneal cavity, and the other through the blood to distal tissues. At a clonal level cells appear to adopt one or the other but not both. This exclusivity may be affected by microenvironmental factors based on the observed differences in blood vs ascites spread between primary tumor sites, and was also influenced by LRRC15 expression. A further understanding of the clonal gene expression patterns could allow the design of targeted therapies to block each of these routes. More immediately, our results suggest that detection of tumor burden in the blood or ascites may be useful as a predictor of probable sites of metastasis in a clinical context. Another interesting example can be found in bowel metastasis. shControl clones present in 8/8 bowel metastasis were shared with primary tumor, proximal metastatic tumor, ascites, lung, heart, both kidneys and contralateral ovary whereas shLRRC15 clones observed in 7/8 sites were exclusive to bowel metastasis only and only clones in 1/8 site was shared with ascites, heart, both kidneys, blood and contralateral ovary. This pattern is consistent with cells metastasizing as oligoclonal clusters as demonstrated previously (27).

The orthotopic implant sites chosen here spatially restricts cells and maximizes co-injected cell competition which may be lost in intravenous or intraperitoneal injections. It is important to remember that our study and others have shown that numbers of cells conventionally implanted to model diseases may not elicit full growth dynamics due to saturating numbers (5). The BC-CIC assay was conducted in an immunodeficient environment which is a potential limitation. The role of immune system in shaping clonal patterns of system-wide metastasis and growth remains to be resolved. Therefore, any barcoding experimental designed to track optimal growth dynamics should carefully consider implanted cell number, site and immune competent status of the recipient animal.

In conclusion, we have described NGS-based cellular DNA barcoding technology and a framework to assist quantitative investigation of system-wide cancer metastasis in preclinical models.

## Materials and Methods

### Construction of barcoded vector libraries

We used pLKO.1 backbone to construct barcode vectors. In brief, we amplified and cloned red shifted Luc gene downstream of MNDU3 promoter and linked via 2A peptide to florescent protein encoding gene which were amplified from Addgene plasmids 48249 (mWasabi), 54569 (mT-sapphire), 54602 (mTag-BFP2) and 74258 (mRuby3). A barcode cloning scaffold was introduced downstream of florescent protein encoding gene. Subsequently, we amplified HA tagged NIS gene, a kind gift from Dr. Stephen Russell (Mayo Clinic) and cloned it downstream of PGK promoter. The barcode linker was created by annealing sense and antisense oligos synthesized by IDT technologies (Table 1). Subsequently, 100μM of sense and antisense oligos were dissolved in T4 ligation buffer and phosphorylated with T4 PNK enzyme (NEB). The mixture was annealed in thermocycler for 5 minutes at 95°C and the annealed oligos were allowed to cool gradually by decreasing temperature to 25°C at 5°C/min. The annealed barcode linker was ligated into the BamHI-Xho1 site of the vector at equimolar ratio. The resulting vector was transformed into MegaX DH10B T1R electrocompetent cells (Invitrogen) using an electroporator (Biorad) and the entire transformed bacteria were inoculated into 1000 ml LB medium containing100 μg/ml carbenicillin (Sigma-Aldrich). After overnight culture, plasmid DNA was extracted with using Maxiprep Plasmid DNA extraction kit (Qiagen). A small aliquot was also inoculated on LB agar plate with 100 g/mL ampicillin to calculate the barcode diversity. Moreover, individual colonies were picked up, plasmids were prepared, and sequence was verified by the Sanger sequencing method.

### HGSOC cancer cell lines

OVCAR5 cells were obtained from Fox Chase Cancer Center, Philadelphia and was maintained using RPMI-1640 media supplemented with 10% FBS and 1% penicillin-streptomycin. LRRC15 knockdown was performed in the OVCAR5 cells with shLRRC15 (Sigma-Aldrich) targeting the 3’UTR [Sequence: GCTATGAAAGAGAGAAGGAAA] using standard transfection guidelines and reagents. Whole cell lysates of OVCAR5-shControl and OVCAR5-shLRRC15 cell lines were subjected to western blot analysis against LRRC15 (ab150376) and GAPDH (sc-47724) antibodies. The blots were visualized using fluorophore-conjugated secondary antibodies (LICOR) and scanned by LI-COR OdysseyFc Imaging System (Nebraska, USA).

### DNA barcoding cancer cell lines

OVCAR5-shControl and OVCAR5-shLRRC15 cell lines were barcoded by lentiviral infection with EGFP_index 1.4 and Ruby3_index 1.2 libraries respectively using 0.8 μg/ml polybrene. After a 24-h incubation, the virus containing medium was removed and cells were cultured for additional 24-h in fresh medium. To ensure that the majority of cells were labeled with a single barcode per cell, for lentiviral infection we used a predetermined dilution of virus (1:500-1:1000) corresponding to infectivity rate <37% based on Poisson statistical modelling. The cells were dissociated using 0.05% trypsin and sorted using stringent gating procedures. The barcoded OVCAR5-shControl (also referred as LRRC15+) and OVCAR5-shLRRC15 (LRRC15-) cells were sorted and mixed in equal numbers (up to 1 million cells) in PBS, loaded on cell injection pipettes and placed on ice until injection. A small portion of sorted cells were verified for purity by FACS analysis. Further, 500-1500 sorted cells were also plated into 6-well plates in 2D clonogeneic assay. After 10 days, colonies were fixed with cold methanol/acetone (1:1) and stained with Giemsa stain. The clonogenic frequency was enumerated and the colony was imaged using Cytation 5 imager and colony area measured using Biotek’s Gen5 software.

### Orthotropic xenotransplantation

The intraoviductal and intraovarian xenotransplant studies in NSG mice were approved by the Mayo Clinic Institutional Animal Care and Use Committee (IACUC). Surgery was performed on one mouse at a time. On the day of surgery, the weight of the mouse was measured and a combination of (90-120 mg/kg) xylazine (10 mg/kg) was administered via intraperitoneal (IP) route. The animal was transferred to cage to induce surgical anesthesia plane. Once, unconscious eye ointment was applied to avoid dryness of eyes and mouse was draped in a cling foil. The mouse was then transferred onto a stage of stereomicroscope platform. Two incision points were marked on the mouse, first between last rib and hip and second between dorsal side and abdomen. The fur was removed using shaver and the skin was scrubbed with povidone-iodine and 70% ethanol. Subsequently, using surgical scissors, a 0.5 cm transversal skin incision was performed at the marked point to expose the fat pad linked to ovary and oviduct. Incision was performed under a stereomicroscope and ovarian fat pad was grabbed in order to minimize the contact with reproductive tract and avoid luteolysis. The fat pad was grasped and held outside the abdominal cavity using a vascular clip. *Oviduct transplantation:*Under the stereomicroscope oviduct (highly coiled structure) was identified and then the ampulla portion of the oviduct that is closer to the ovary was located. The ampulla portion was then placed horizontally on the surgical platform and a small transversal cut nearly three fourth of the circumference on the distal part of ampulla using a micro-scissor. The glass needle was slowly inserted through the cut site and cell suspension in trypan blue buffer was injected. Few air bubbles were also injected to indicate successful injection (*see* Figure S4A-B). *Ovary transplantation:* The ovary has a characteristic reddish appearance under the stereomicroscope. Ovary was held using forceps. Subsequently, cell injection pipette was inserted into the bursa of ovary and cells were injected. After the injection, the glass capillary was slowly withdrawn, and tension was exerted by forceps and grasp was released. The ovary and oviduct put back into the abdominal cavity by grabbing the fat pad using forceps. The muscle and skin layer were then sutured using 5/0 absorbable sutures. Children’s ibuprofen (40 mg/kg, 0.2 mg/ml of water) was given as analgesic in drinking water 48 hours before and after surgical procedure.

### Bioluminescence imaging

The tumor growth and volume were non-invasively monitored using Xenogen IVIS optical imaging system. Briefly, animals were anesthetized using 2% isoflurane in an induction chamber and injected with 0.2 ml (150 mg/kg) of luciferin substrate through intra-peritoneal route using 25-gauge needles. Subsequently, the animals were placed on a pre-warmed stage in IVIS imaging chamber and 2% isoflurane anesthesia was maintained during the course of procedure via nose cone. Living Image^®^ Software from Xenogen was used for image acquisition. The animals were returned to respective cages following the procedure and observed for recovery.

### Barcode amplification

At the endpoint primary tumor and other organs were harvested and snap frozen. Genomic DNA was extracted from the frozen tissues (10 mg) with a MasterPure Complete DNA and RNA purification Kit (Lucigen). We used Q5^®^High-Fidelity 2X Master Mix (NEB) to amplify the barcode sequence for NGS by introducing Illumina adaptors and 5-bp-long index sequences. The sampling of sufficient template coverage was ensured by parallel PCR reactions. For each PCR reaction, 1 μg of genomic DNA was spiked with “spike in” controls and used as a template. The sequence information for the primers used for barcode amplification can be found in Supplementary Table 2. Labeling each sample with 1 of the 20 unique indices enabled us to multiplex and sequence up to 20 samples at once.

### NGS barcode sequencing

Completed libraries were spiked with 10% commercially prepared PhiX library (Illumina) to increase base diversity for improved sequencing quality. Samples were sequenced at 1 sample per lane to generate approximately 85 to 95 million total reads per sample. The single read flow cells were sequenced as 100 base single end reads on an Illumina HiSeq 2500 using TruSeq Rapid SBS sequencing kit version 1 and HCS version 2.0.12.0 data collection software. Base-calling is performed using Illumina’s RTA version 1.17.21.3.

### Barcode data processing and threshold determination

In the preprocessing step, the barcode sequences were extracted from the raw FASTQ files based on the sequences of index 1 and index 2 at either ends of the amplicon. The number of sequences sharing the same unique barcode were then counted and barcodes from the same library with no more than 2 mismatched base pairs were merged as these likely arose due to sequencing errors (5, 11). A barcode count table was generated with respect to all identified clones with unique index 1, index 2 and barcode sequence combination. Raw counts were then normalized to fractional read values (FRVs) by dividing the raw clone counts by the total sample size: FRV =raw Counts/total raw counts in sample.

The CIC Calculator filtered the noise clones using the dedicated spike-in index 1.6 (GTCA) library in each sample as a ground truth. To do so, the true positive rate (TPR) and false positive rate (FPR) were calculated as below.

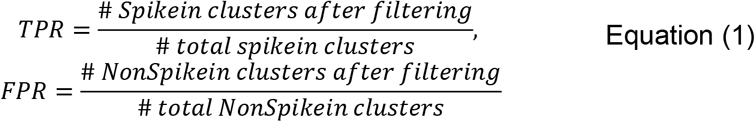

The fractional read value (FRV) threshold was then determined based on the optimal accuracy from the receiver operating characteristic (ROC) curves. This was done either using the local spike-in controls (those from each individual sample), or based on a pool of spike-in controls from all samples (global). For our downstream analyses, the global setting was used. Small barcode clones with FRV lower than the threshold were filtered out. Next a linear fit was performed between the log10(FRVs) of the spike-in controls and the known input copy numbers (ignoring single copy inputs). The resulting fit was then used to convert FRVs into absolute cell counts which could be used in subsequent analyses.

### Analyses of clonal patterns

All unique clones appearing above detection threshold were first identified for each mouse and all sites with detected values for each identified. As we know that all remaining sites these clones were below the threshold of detection, and many identical values can interfere with various tests, we interpolated values for each unique clone at below threshold sites. To do so we fit a left-censored Poisson distribution to each site for each mouse based on observed clones at that site. Lambda values between 0 and 100 were tested in increments of 0.1 and the value selected which minimized the sum of squared error. From this lambda value, we could then infer the expected frequency of cells at below threshold values. Clones were then assigned random cell numbers below thresholds with the probability based on the Poisson fit (i.e., filled in the below threshold values of the best-fit Poisson distribution). For those sites that lacked sequencing data due to a failure to amplify of the barcode, we once again know that they must have below threshold levels. As no distribution could be fit, a uniform random distribution was used to assign below threshold values in these cases. Missing data for which tissues had not been collected were left as missing. These interpolated cell number matrices were used for subsequent analyses. All analyses and statistics for this and following sections were performed in R (version 3.6.1).

To determine whether there was a relationship between metastatic sites, we first calculated pair-wise Spearman’s correlations between sites for each mouse based on the detected cell numbers at each metastatic site. These matrices were then converted to distance matrices based on the cosine theorem: 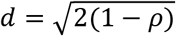. As an initial measure of whether these mice showed similar patterns (and thus it was reasonable to combine distance matrices), we performed a congruence among distance matrices test using the R package ‘ape’ (version 5.4.1). We next took a mean of the distance matrices across mice ignoring missing values to get the average pattern of relationships. This was also done for only intraovarian injections, and for intraoviductal mice only, including only those sites which were measured in all mice in that set. Each of these methods was used to perform a classical multi-dimensional scaling (MDS). In order to obtain a measure of the biological variability of these patterns, we first calculated a principal component analysis (PCA) on the mean distance matrix (approximately equivalent MDS). The PCAs were then used to embed the matrices from each individual mouse into the same PCA space.

Given the apparent association of blood and ascites to specific other metastatic sites, we next set out to estimate the relative importance of blood and ascites to metastasis to each site. To do so we next fit a series of multivariate linear regression models. In each case, a model was designed to predict one site given the cell numbers in Blood, Ascites, and the Primary Tumor as predictors: 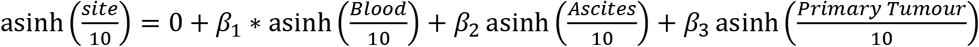. Clones with missing measurements in any predictor or in the site being predicted were excluded from that given model. Adjusted R^2^ and p-values were calculated for each fit, and false-discovery rate was then calculated across all models/predictors to correct for multiple testing.

In order to identify common patterns of metastatic spread we next performed clustering on the overall dataset. This was done in several stages. First, we removed all clones which had less than 10 cells detected at all sites and assigned these to a Low (“L”) cluster. From the remaining clones, we randomly sampled 100 of these from each mice and calculated a pair-wise Pearson’s correlation between the asinh(cell #/10) of each cell. Where measurements were missing, the correlation was calculated between those cells including only the values measured in both clones (ie. pair-wise complete measurements). A k-nearest neighbor (KNN) graph was then constructed based with a maximum of 10 nearest neighbors per cell and including edges only if the Pearson’s correlation was 0.9 or greater using the R packages ‘igraph’ (version 1.2.6). The Leiden clustering algorithm (28) was then run on the KNN graph with a resolution parameter of 0.5 to generate an initial set of clusters using the R package ‘leiden’ (version 0.3.6). Mean values for each site were then calculated for each cluster and cells which were not part of the initial clustering assigned to their most correlated cluster mean (Pair-wise complete Pearson’s correlation of the asinh(cell #/10)) if that most correlated cluster was at least 0.7. If not, these cells were left unassigned. Next, all clusters where were only detected in a single mouse and those with 10 clones or less were removed to eliminate possible artefacts. Finally, to minimize over-clustering, clusters with cluster means with a Pearson’s correlation of 0.7 or more were merged. This resulted in a total of 17 clusters, a cluster of clones which were borderline detectable in all sites, and 295 unassigned clones. As a visual assessment of cluster relationships and cluster solution success, dimensionality reduction was performed on a distance matrix calculated from the pair-wise complete Pearson’s correlation matrix of asinh(cell #/10) between all clones (converted as: 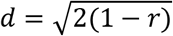). This was done both by classical MDS, as well as by UMAP using the R package ‘umap’ (version 0.2.7.0).

To identify the effects of LRRC15 and injection site on clone numbers both overall and for each cluster, a series of Aligned Rank Transform (ART) 2-factor ANOVAs were performed (29) using the R package ‘ARTool’ (version 0.10.8). In all cases clone number was used as response, while genotype (LRRC15 wild-type vs shRNA) and injection site (intraovarian vs intraoviductal) and the possible interaction between them used as predictors. For comparisons of clone numbers across clusters, p-values were multiple testing corrected by false discovery rate (FDR). As blood was not measured in most intraoviductal mice but was a very strong predictor of engraftment at distant metastatic sites, we performed an elastic net (EN) logistic regression to predict whether a clone would be present in the blood or not. This model used asinh (cell #/10) for all sites measured in all mice (other than blood and ascites) as predictors for the binary variable of detectable or not in blood. To implement this model, we first split the data into training and validation sets, with 70% of the clones present in the blood from each of the intraovarian mice included in the training set, and double that number of clones which were not present in the blood from those mice. This over-representation of blood clones was done to ensure that a sufficient number of positives were included in the training data and prevent specificity (as most clones were not in blood) from being the driving force of the scoring. The remaining 30% of blood clones and the remaining non-blood clones were kept back as a validation set. We then performed hyperparameter fitting with a fixed alpha value of 0.5 (to balance between model sparsity and grouping) with varying lambda and gamma regularization penalties using 5-fold cross validation with area under the receiver-operator curve (AUC) as the scoring parameter using the R package ‘glmnet’ (version 4.0.2). Overall prediction accuracy, sensitivity, and specificity was then scored on a per-mouse basis in the training and validation sets. Another model was then generated to predict whether a clone was present in ascites or not. In this case, the training set included 70% of clones present in ascites from the intraovarian mice and an equal number (as ascites clones were more frequent than blood) which were not present in ascites. Hyperparameter optimization, training and validation was then performed as was done for blood.

## Supplementary data

**Figure S1**. **Cellular DNA barcoding libraries. A)** illustration of the barcode library vector design showing MNDU3 promoter driven expression of red-shifted luciferase (Luc) gene fused to fluorescent protein (FP) gene (EGFP, mWasabi, mRuby3, T-Sapphire and mTag-BFP2) via a 2A linker peptide. A semi-random barcode sequence is cloned downstream of FP gene and 5’ end of barcode sequence is flanked by one of the six indices (index 1.1-1.6) that distinguish the libraries. PGK promoter drives the expression of hemagglutinin (HA)-tagged human sodium-iodine symporter (NIS) gene. **B)** estimated diversities and lentivirus titer of 25 high complexity multi-functional DNA barcode libraries. **C)** functional validation of Luc reporter gene. Plot show photon emission over a period of 30 minutes in cells expressing barcode library vector encoded Luc gene compared to control cells. **D)** FACS analysis demonstrating the multiplexing capabilities of mRuby3, Tag-BFP2 mWasabi, and T-Sapphire FPs. **F)** functional analysis of NIS gene. Plot shows in vitro ^[18]^TFB uptake by HA-tagged hNIS reporter expressing cells and control cells.

**Figure S2. Spike-in control barcode library and multiplexed library construction. A)** table showing copy numbers, barcode and index sequences of plasmids used to create “spike-in” control library for NGS sequencing. **B)** illustration showing multiplexed library preparation steps for NGS sequencing.

**Figure S3. FACS purification and in vitro testing. A)** representative FACS plots of sorted cells. **B)** purities of sorted cells analyzed by FACS. **C)** representative clonogenic assay dish after Giemsa staining showing OVCAR5-shControl and shLRRC15 colonies.

**Figure S4. Orthotopic xenotransplantation. A)** picture showing the glass needle (130-150μm diameter) that was used for co-transplantation of cells at intraoviductal or intraovarian sites. **B)** micrographs and schematics of steps involved in intraoviductal and intraovarian xenotransplant in NSG mice.

**Figure S5. Necropsy.** Images showing visceral anatomy, primary tumors and metastasis to various organs in NSG mice transplanted at intraoviductal (n=10) and intraovarian (n=10) sites. The images demonstrate 100% take rate for co-transplanted cells.

**Figure S6. Characteristics of tumors. A-H)** tissue/organ weights. Groups were compared using student t-test. **I)** correlation between injected cell dose and tumor weight.

**Figure S7. Molecular autopsy.** PCR analysis of primary tumor and various organs harvested at endpoint for the presence of barcoded cells across anatomical sites.

**Figure S8. Largest clones across sites**. Plots showing significant difference in the estimated size of the largest clones in primary tumors and organs harvested from intraoviductal **(A)** and intraovarian **(B)** cell injection sites. LRRC15- and LRRC15+ groups were compared using paired two-tailed t-test, ***<0.0005 and ****<0.0001.

**Figure S9. CIC-MM plots depicting system-wide metastasis in intraoviductal co-injected mice.**

**Figure S10. CIC-MM plots depicting system-wide metastasis in intraovarian co-injected mice.**

**Figure S11. Metastatic site relationships by tumor primary site. A,B)** average relationships between sites of clone growth for intraovarian only **A)** and intraoviductal only **B)**. Sites of growth are shown embedded by classical multi-dimensional scaling (MDS) using the averaged distance-transformed Spearman’s correlation across mice. Metastatic site is indicated on the plot, and the relative distance between sites is inversely proportional to the strength of their correlation (ie, the closer on the plot they appear, the more strongly correlated). **C,D)** per-mouse relationship of clone growth including only intraovarian only **C)** and intraoviductal only **D)** the plot shows a PCA embedding based on the averaged distance matrix across mice (approximately equivalent to MDS) followed by embedding the per-mouse relationships into this space as a measure of how consistent these relationships are across individual animals. Injection site is indicated by point shape and site of growth is indicated by colour as shown in the legend. **E)** coefficients of are shown for a series of multivariate linear regression models including only mice injected at the intraovarian site using clone size detected in the blood, ascites, and primary tumor to predict levels in each other site (all asinh(x/10) transformed to maintain linearity). Higher coefficients (values on the y-axis) represent a greater predictive power of that site for the overall clone size at the indicated site. Lines showing 2x the standard error are shown for each coefficient.

**Figure S12. Relationships between each site, blood, ascites, and primary tumor size.** Plots show asinh(x/10) scaled cell numbers per clone of each site on the y-axis and blood (left), ascites (middle) or primary tumor site (right) on the x-axis. Clones from all mice are shown in **A)**, only clones from mice injected at the Intraovarian site in **B)**, and only clones from mice injected at the intraoviductal site in **C)**. grey boxes show detection thresholds. Any clone with a missing value in either of the two displayed axes for each plot is not included.

**Figure S13. Cluster patterns all clusters**. Clonal engraftment patterns are shown for all. Clusters L, I, IV, VIII and X are reproduced here for completeness from Figure X3. Lines connect cell numbers detected for each clone between sites. Missing values for a given clone are shown as gaps between adjacent line segments. Clone numbers are asinh(cell #/10) transformed with the linear numbers indicated. The total number of clones which were part of a cluster, and the number of mice with at least one clone that was a member of that cluster for each injection site are shown in the top left. The cluster identifier for each is shown in the top right.

**Figure S14. Clone numbers by injection site and LRRC15 status**. The number of detectable clones which are part of each cluster is shown for each mouse separated by injection site and LRRC15 status (“-” = LRRC15 shRNA, “+” = LRRC15 wild-type). Lines indicate median values. The cluster identifier for each is shown in the top right. Clusters L, I, IV, VIII and X are reproduced here for completeness from Figure X3.

**Figure S15. Metastatic site usage predictions. A)** hyperparameter fitting for the blood EN model. Cross-validation area-under-the-curve (AUC) is indicated for a range of values of the regularization parameter lambda for 4 values of the regularization parameter gamma. The first dotted line indicates the hyperparameter values which maximize AUC, and the second the most regularized values (highest lambda and gamma) which are within 1 standard error of the maximum AUC. The numbers at the top of the plot show the number of non-zero predictors included in the model. **B)** percentage of clones per mouse with measured detection in blood separated by injection site. **C)** hyperparameter fitting for the ascites EN model (as for **A**). **D)** overall accuracy (left), sensitivity (middle) and specificity (right) of an elastic net (EN) logistic regression to predict whether a given clone is present (detectable) in ascites. The EN model was trained on 70% of ascites detectable clones from each mouse in the intraovarian cohort and an equal number of non-ascites clones with remaining clones kept back for the validation set. Each point represents the predictions for clones from one mouse. Each plot shows the relevant metric for the training and validation sets. **E)** the coefficients retained for each site in the final EN model for predicting whether a clone will be detectable in ascites or not and their magnitude. **F)** percentage of clones per mouse predicted to be detectable in ascites by the EN model separated by injection site.

**Table S1. Sensitivity data for ‘spike in’ internal calibration controls.**

**Table S2. Number of cells co-injected in intraoviductal and intraovarian injected mice.**

**Table S3. Percentage of CIC classes in intraoviductal and intraovarian injected mice.**

**Table S4. Estimated maximum clone size per organ in intraoviductal and intraovarian injected mice.**

**Table S5. Hexcodes for MM plots**.

**Table S6. Linear model statistics.** statistics for linear models. “Model” contains the growth site for which the predictors are predicting the value. “Predictors” has for a given row, which predictor (Ascites, Blood, or Primary Tumour) the values pertain to. “Estimate” shows the estimated coefficient from the linear model on the scale asinh(Cell Number/10). “Standard Error” contains the standard error of the estimate. “t.value”has the raw t statistic. “P-value” has unadjusted p values based on the t statistic and “False Discovery Rate” contains the false discovery rate (multiple testing adjusted measure of significance).

**Table S7. Significance**. ART ANOVA statistics for each cluster. “Cluster” contains the cluster for a given ART ANOVA. “Term” contains the variable including injection site (“Site”), LRRC15 status (“Genotype”) or the interaction between the two (“Site:Genotype”). “p” contains the raw p values for each Term, and “FDR” contains the false-discovery rate (multiple testing adjusted measure of significance).

**Video S1. Video demonstrating clonal patterns of metastasis in mouse #67.** Each frame represents an individual clone and their system-wide linkages. All clones in mouse #67 are shown.

## Author contributions

SMA and JS conducted the experiments. JBG and MS helped develop surgical method. XT, DK, KK and NK analyzed the results, performed statistical analysis. UR and VS provided the cancer cell lines. NK conceptualized and designed the study, interpreted the data, and wrote the manuscript. NK supervised the study. All authors contributed to the drafting of the manuscript.

## Acknowledgements

JS received travel fellowship from Beijing Lihuanying Medical Foundation, China to visit NK lab. DK is supported through a Fonds de recherche du Québec – Santé (FRQS) Junior 1 salary award, and partial funding for this work were provided by generous donations from the Fondation Marcelle et Jean Coutu through the IRIC philanthropic funds. The study was supported by research award received by NK from the Mayo Clinic’s Department of Laboratory Medicine and Pathology (DLMP).

## References

1. Riggi N, Aguet M, Stamenkovic I. Cancer Metastasis: A Reappraisal of Its Underlying Mechanisms and Their Relevance to Treatment. Annu Rev Pathol. 2018;13:117–40.

2. Fares J, Fares MY, Khachfe HH, Salhab HA, Fares Y. Molecular principles of metastasis: a hallmark of cancer revisited. Signal Transduct Target Ther. 2020;5(1):28.

3. Lan X, Jorg DJ, Cavalli FMG, Richards LM, Nguyen LV, Vanner RJ, et al. Fate mapping of human glioblastoma reveals an invariant stem cell hierarchy. Nature. 2017;549(7671):227–32.

4. Nguyen LV, Cox CL, Eirew P, Knapp DJ, Pellacani D, Kannan N, et al. DNA barcoding reveals diverse growth kinetics of human breast tumour subclones in serially passaged xenografts. Nat Commun. 2014;5:5871.

5. Nguyen LV, Makarem M, Carles A, Moksa M, Kannan N, Pandoh P, et al. Clonal analysis via barcoding reveals diverse growth and differentiation of transplanted mouse and human mammary stem cells. Cell Stem Cell. 2014;14(2):253–63.

6. Nguyen LV, Pellacani D, Lefort S, Kannan N, Osako T, Makarem M, et al. Barcoding reveals complex clonal dynamics of de novo transformed human mammary cells. Nature. 2015;528(7581):267–71.

7. Bhang HE, Ruddy DA, Krishnamurthy Radhakrishna V, Caushi JX, Zhao R, Hims MM, et al. Studying clonal dynamics in response to cancer therapy using high-complexity barcoding. Nat Med. 2015;21(5):440–8.

8. Hata AN, Niederst MJ, Archibald HL, Gomez-Caraballo M, Siddiqui FM, Mulvey HE, et al. Tumor cells can follow distinct evolutionary paths to become resistant to epidermal growth factor receptor inhibition. Nat Med. 2016;22(3):262–9.

9. Kebschull JM, Zador AM. Cellular barcoding: lineage tracing, screening and beyond. Nat Methods. 2018;15(11):871–9.

10. Gerrits A, Dykstra B, Kalmykowa OJ, Klauke K, Verovskaya E, Broekhuis MJ, et al. Cellular barcoding tool for clonal analysis in the hematopoietic system. Blood. 2010;115(13):2610–8.

11. Cheung AM, Nguyen LV, Carles A, Beer P, Miller PH, Knapp DJ, et al. Analysis of the clonal growth and differentiation dynamics of primitive barcoded human cord blood cells in NSG mice. Blood. 2013;122(18):3129–37.

12. Naik SH, Schumacher TN, Perie L. Cellular barcoding: a technical appraisal. Exp Hematol. 2014;42(8):598–608.

13. Merino D, Weber TS, Serrano A, Vaillant F, Liu K, Pal B, et al. Barcoding reveals complex clonal behavior in patient-derived xenografts of metastatic triple negative breast cancer. Nat Commun. 2019;10(1):766.

14. Seth S, Li CY, Ho IL, Corti D, Loponte S, Sapio L, et al. Pre-existing Functional Heterogeneity of Tumorigenic Compartment as the Origin of Chemoresistance in Pancreatic Tumors. Cell Rep. 2019;26(6):1518–32 e9.

15. Porter SN, Baker LC, Mittelman D, Porteus MH. Lentiviral and targeted cellular barcoding reveals ongoing clonal dynamics of cell lines in vitro and in vivo. Genome Biol. 2014;15(5):R75.

16. Echeverria GV, Powell E, Seth S, Ge Z, Carugo A, Bristow C, et al. High-resolution clonal mapping of multi-organ metastasis in triple negative breast cancer. Nat Commun. 2018;9(1):5079.

17. Wagenblast E, Soto M, Gutierrez-Angel S, Hartl CA, Gable AL, Maceli AR, et al. A model of breast cancer heterogeneity reveals vascular mimicry as a driver of metastasis. Nature. 2015;520(7547):358–62.

18. Roh V, Abramowski P, Hiou-Feige A, Cornils K, Rivals JP, Zougman A, et al. Cellular Barcoding Identifies Clonal Substitution as a Hallmark of Local Recurrence in a Surgical Model of Head and Neck Squamous Cell Carcinoma. Cell Rep. 2018;25(8):2208–22 e7.

19. Echeverria GV, Ge Z, Seth S, Zhang X, Jeter-Jones S, Zhou X, et al. Resistance to neoadjuvant chemotherapy in triple-negative breast cancer mediated by a reversible drug-tolerant state. Sci Transl Med. 2019;11(488).

20. Aalam SMM, Beer PA, Kannan N. Assays for functionally defined normal and malignant mammary stem cells. Adv Cancer Res. 2019;141:129–74.

21. Kokkaliaris KD, Lucas D, Beerman I, Kent DG, Perie L. Understanding hematopoiesis from a single-cell standpoint. Exp Hematol. 2016;44(6):447–50.

22. Mariani A, Wang C, Oberg AL, Riska SM, Torres M, Kumka J, et al. Genes associated with bowel metastases in ovarian cancer. Gynecol Oncol. 2019;154(3):495–504.

23. Bermejo-Alvarez P, Park KE, Telugu BP. Utero-tubal embryo transfer and vasectomy in the mouse model. J Vis Exp. 2014(84):e51214.

24. Bast RC, Jr., Hennessy B, Mills GB. The biology of ovarian cancer: new opportunities for translation. Nat Rev Cancer. 2009;9(6):415–28.

25. Talmadge JE, Wolman SR, Fidler IJ. Evidence for the clonal origin of spontaneous metastases. Science. 1982;217(4557):361–3.

26. Nicolson GL, Custead SE. Tumor metastasis is not due to adaptation of cells to a new organ environment. Science. 1982;215(4529):176–8.

27. Aceto N, Bardia A, Miyamoto DT, Donaldson MC, Wittner BS, Spencer JA, et al. Circulating tumor cell clusters are oligoclonal precursors of breast cancer metastasis. Cell. 2014;158(5):1110–22.

28. Traag VA, Waltman L, van Eck NJ. From Louvain to Leiden: guaranteeing well-connected communities. Sci Rep. 2019;9(1):5233.

29. Wobbrock J.O. FL, Gergle D., Higgins J.,. The Aligned Rank Transform for Nonparametric Factorial Analyses Using Only ANOVA Procedures. 2011. p. 143–6.

