## Supplementary material for "Barcoded Competitive Clone-Initiating Cell (BC-CIC) Analysis Reveals Differences in Ovarian Cancer Cell Genotype and Niche Specific Clonal Fitness During Growth and Metastasis In Vivo": Figure S1

**A**

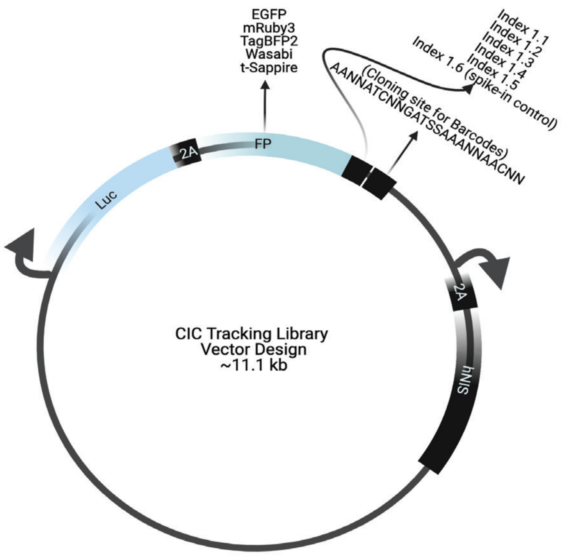


**B**

| Library Number | Description | Estimated diversity | Transduction units/ml |
|---|---|---|---|
| L1 | EGFP_Barcode _Index_1 | $1.1 X 10^6$ | $3.98 x 10^{12}$ |
| L2 | EGFP_Barcode _Index_2 | $1.3 X 10^6$ | $2.07 X 10^{13}$ |
| L3 | EGFP_Barcode _ Index_3 | $2.1 X 10^6$ | $1.84 X 10^{13}$ |
| L4 | EGFP_Barcode _ Index_4 | $1.6 X 10^6$ | $1.2 X 10^{13}$ |
| L5 | EGFP_Barcode _ Index_5 | $2.2 X 10^6$ | $2.32 X 10^{13}$ |
| L6 | Ruby3_Barcode _Index_1 | $3.0 X 10^6$ | $1.3 X 10^{13}$ |
| L7 | Ruby3_Barcode _Index_2 | $3.0 X 10^6$ | $1.72 X 10^{13}$ |
| L8 | Ruby3_Barcode _ Index_3 | $2.8 X 10^6$ | $2.0 X 10^{13}$ |
| L9 | Ruby3_Barcode _ Index_4 | $2.0 X 10^6$ | $1.86 X 10^{13}$ |
| L10 | Ruby3_Barcode _ Index_5 | $2.1 X 10^6$ | $2.1 X 10^{13}$ |
| L11 | TagBFP2_Barcode _Index_1 | $2.4 X 10^6$ | $2.8 x 10^{13}$ |
| L12 | TagBFP2_Barcode _Index_2 | $2.3 X 10^6$ | $2.32 X 10^{13}$ |
| L13 | TagBFP2_Barcode _Index_3 | $2.4 X 10^6$ | $1.76 X 10^{13}$ |
| L14 | TagBFP2_Barcode _Index_4 | $2.1 X 10^6$ | $2.34 X 10^{13}$ |
| L15 | TagBFP2_Barcode _Index_5 | $2.3 X 10^6$ | $2.1 X 10^{13}$ |
| L16 | Wasabi_Barcode _Index_1 | $3.0 X 10^6$ | $1.8 X 10^{13}$ |
| L17 | Wasabi_Barcode _Index_2 | $2.3 X 10^6$ | $1.8 X 10^{13}$ |
| L18 | Wasabi_Barcode _ Index_3 | $2.8 X 10^6$ | $2.1 X 10^{13}$ |
| L19 | Wasabi_Barcode _ Index_4 | $2.4 X 10^6$ | $2.24 X 10^{13}$ |
| L20 | Wasabi_Barcode _ Index_5 | $1.8 X 10^6$ | $2.1 X 10^{13}$ |
| L21 | Sapphire_Barcode _Index_1 | $2.0 X 10^6$ | $2.12 X 10^{13}$ |
| L22 | Sapphire_Barcode _Index_2 | $1.8 X 10^6$ | $2.0 X 10^{13}$ |
| L23 | Sapphire_Barcode _ Index_3 | $2.1 X 10^6$ | $2.3 X 10^{13}$ |
| L24 | Sapphire_Barcode _ Index_4 | $1.3 X 10^6$ | $2.3 X 10^{13}$ |
| L25 | Sapphire_Barcode _ Index_5 | $1.8 X 10^6$ | $1.9 X 10^{13}$ |

**C**

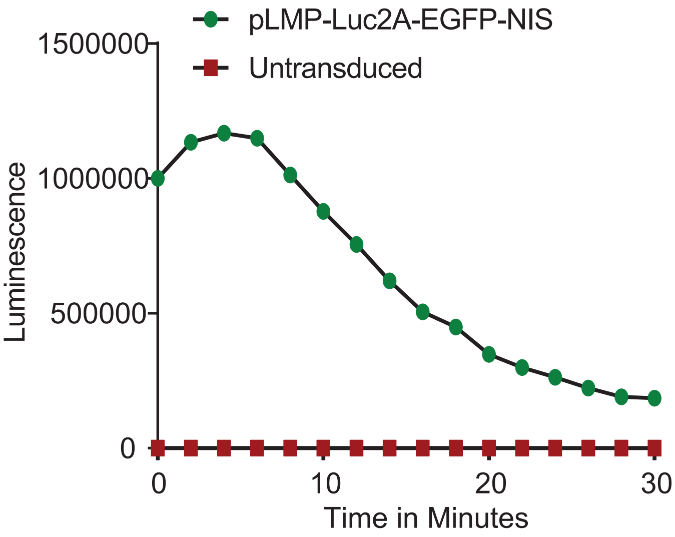


**D**

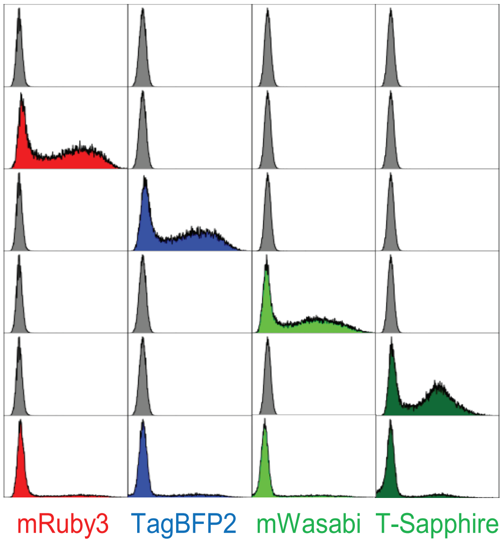


**E**

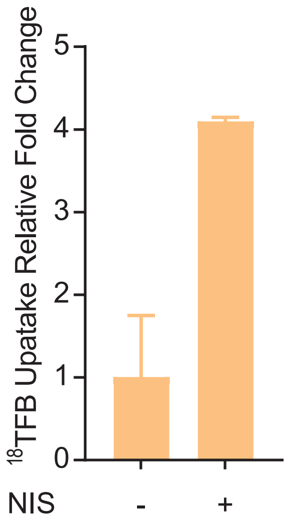
