## Supplementary material for "Barcoded Competitive Clone-Initiating Cell (BC-CIC) Analysis Reveals Differences in Ovarian Cancer Cell Genotype and Niche Specific Clonal Fitness During Growth and Metastasis In Vivo": Figure S2

**Supplementary Figure 2**

**A**

| Plasmid No | Expected Copies | Index | Spike in Barcode Sequence |
|---|---|---|---|
| 1 | 10K | GTCA | AAGTAA CA ATC GT GAT CG AAA TG GGT CG AAC TT AGCA |
| 2 | 1K | GTCA | AAGTAA GA ATC GA GAT CC AAA AG GGT TA AAC AA AGCA |
| 6 | 1K | GTCA | AAGTAA AA ATC GG GAT GC AAA AC GGT TT AAC GT AGCA |
| 7 | 1K | GTCA | AAGTAA GG ATC GT GAT CG AAA TC GGT AT AAC CT AGCA |
| 10 | 100 | GTCA | AAGTAA GG ATC GT GAT CG AAA AT GGT CC AAC TT AGCA |
| 11 | 100 | GTCA | AAGTAA TT ATC AC GAT GC AAA TG GGT AT AAC AT AGCA |
| 13 | 100 | GTCA | AAGTAA GT ATC AT GAT GG AAA AT GGT TT AAC AT AGCA |
| 16 | 10 | GTCA | AAGTAA GG ATC GC GAT GG AAA TG GGT AG AAC CC AGCA |
| 17 | 10 | GTCA | AAGTAA TA ATC GG GAT GC AAA GG GGT GT AAC GG AGCA |
| 26 | 10 | GTCA | AAGTAA GT ATC AG GAT CG AAA CT GGT GG AAC GA AGCA |
| 27 | 10 | GTCA | AAGTAA TA ATC CT GAT GG AAA GT GGT GG AAC TG AGCA |
| 30 | 10 | GTCA | AAGTAA GA ATC TA GAT CG AAA TG GGT GG AAC GG AGCA |
| 33 | 1 | GTCA | AAGTAA TT ATC GC GAT GC AAA AC GGT AG AAC GG AGCA |
| 34 | 1 | GTCA | AAGTAA CT ATC AG GAT GG AAA CA GGT GG AAC GA AGCA |
| 35 | 1 | GTCA | AAGTAA GG ATC GA GAT CG AAA TC GGT AT AAC TA AGCA |
| 36 | 1 | GTCA | AAGTAA CC ATC AT GAT CC AAA TG GGT TT AAC TA AGCA |
| 37 | 1 | GTCA | AAGTAA AC ATC AC GAT GG AAA TC GGT AA AAC GT AGCA |

**B**

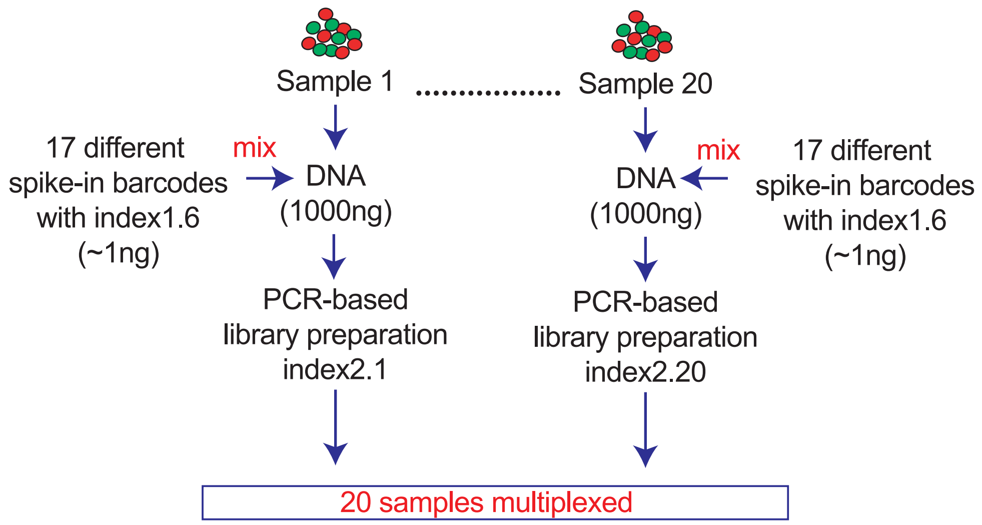
