## Supplementary figures and images for "Barcoded Competitive Clone-Initiating Cell (BC-CIC) Analysis Reveals Differences in Ovarian Cancer Cell Genotype and Niche Specific Clonal Fitness During Growth and Metastasis In Vivo"

### Figure S3

**Figure S3**

**A**

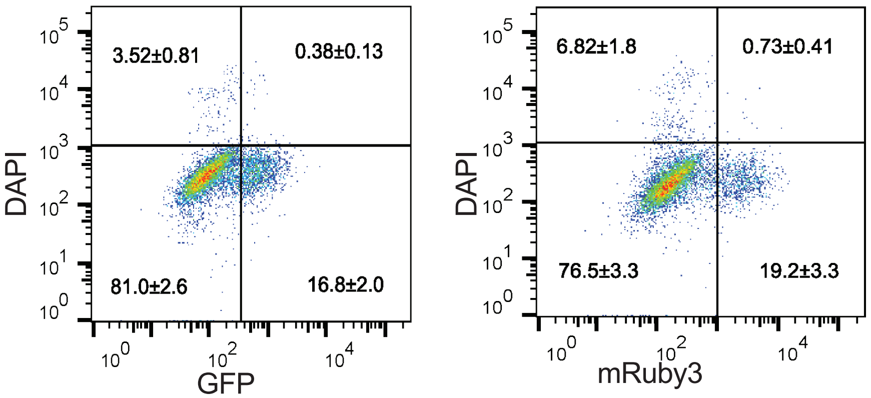


**B**

| Post-sort analysis | | |
|---|---|---|
| Sort# | %GFP | %Ruby |
| 1 | 85.4 | 72.5 |
| 2 | 46.1 | 60.5 |
| 3 | 55.6 | 72.8 |
| 4 | 63.3 | 68 |
| 5 | 62.9 | 58.1 |
| 6 | 71.6 | 61 |
| SEM | 65.3±2.7 | 64.1±5.5 |

**C**

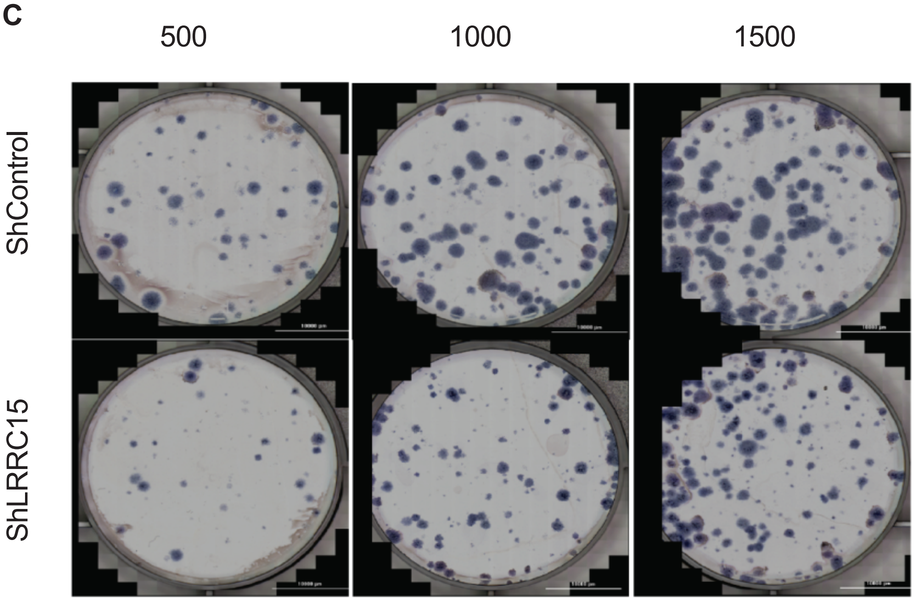

### Figure S4

**Figure S4**

**A**

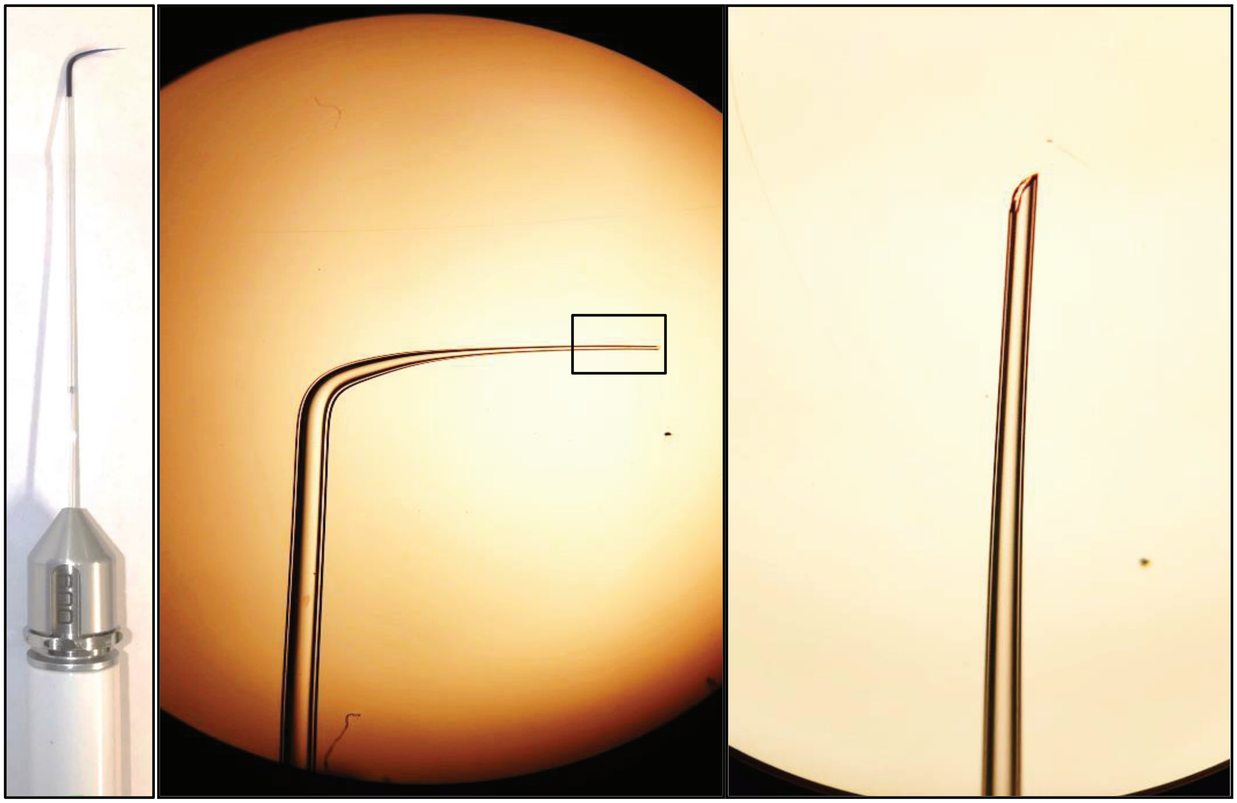

**B**

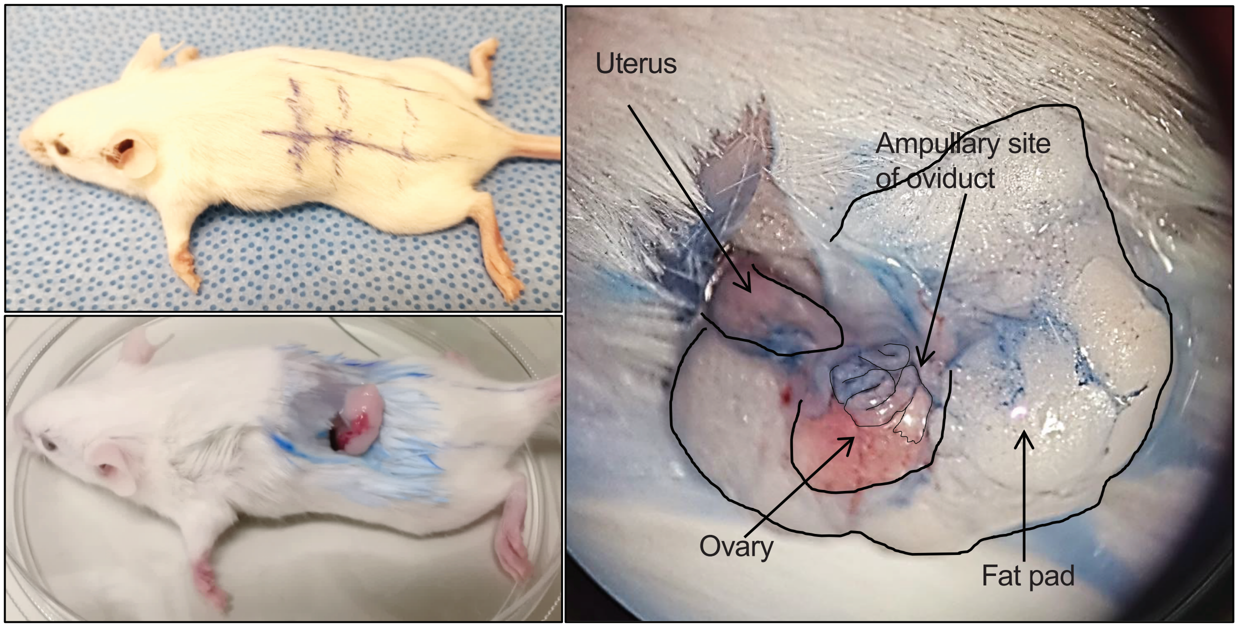

### Figure S5

**Supplementary Figure 5**

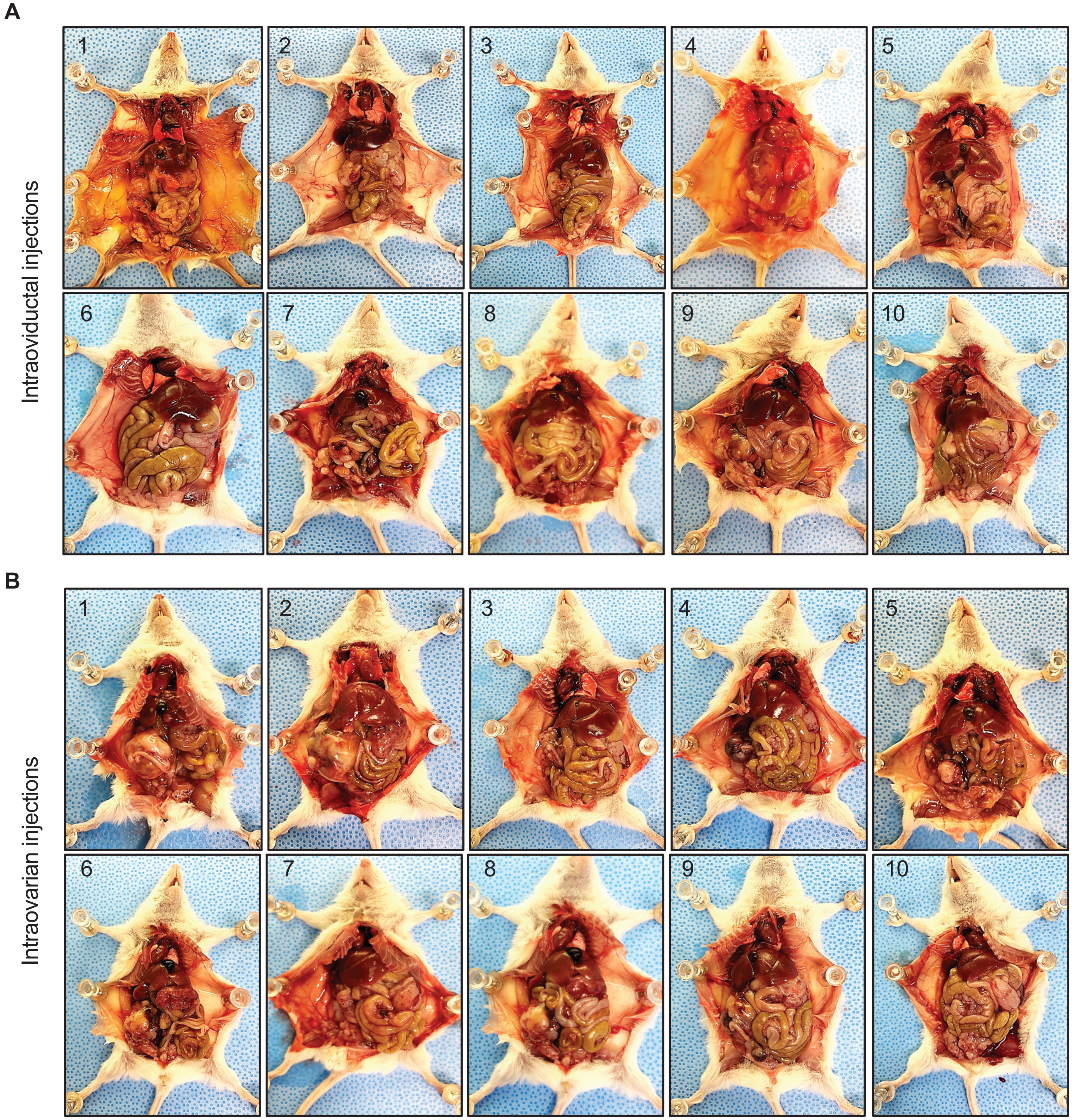

### Figure S6

**Supplementary Figure 6**

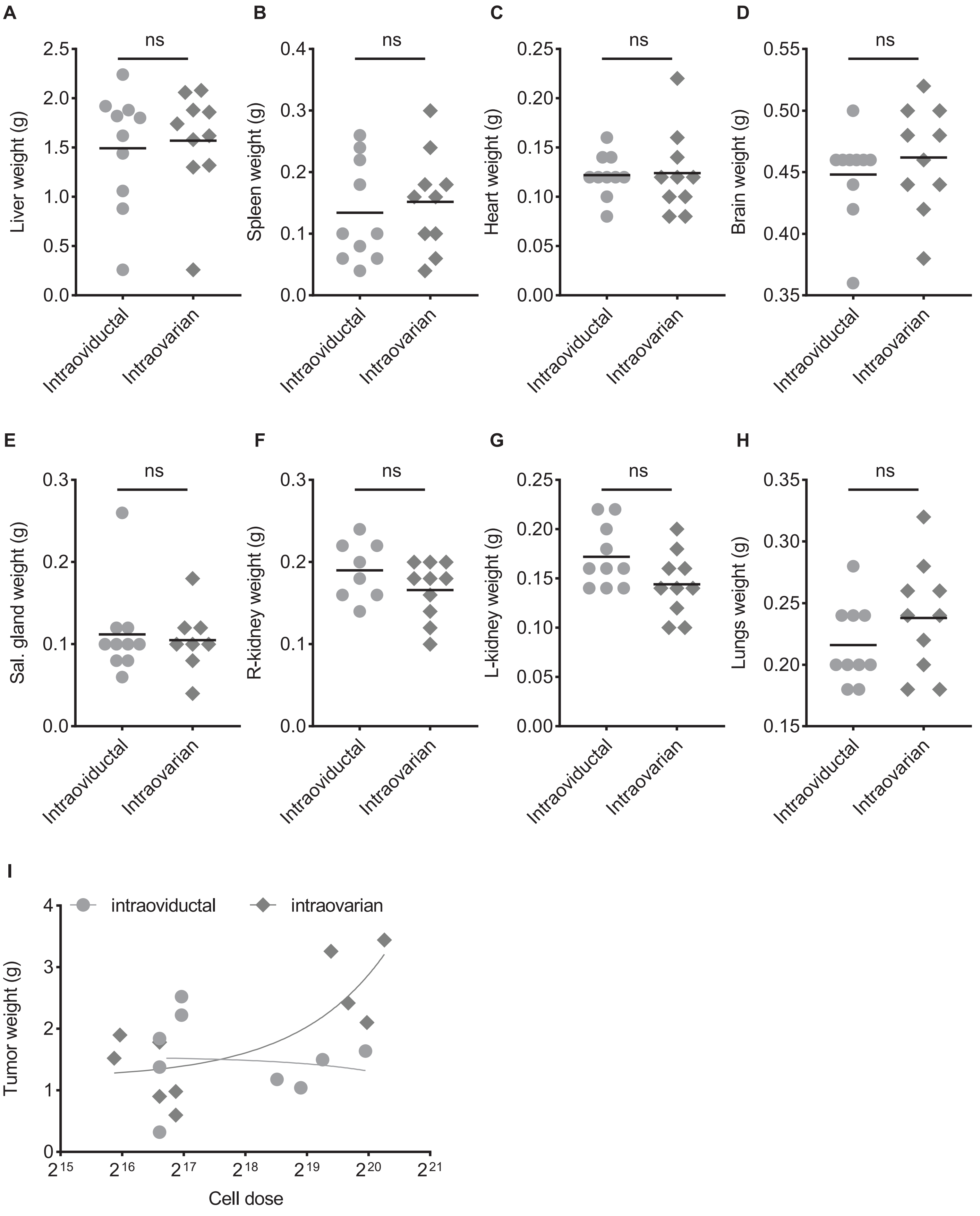

### Figure S7

#### Supplementary Figure 7

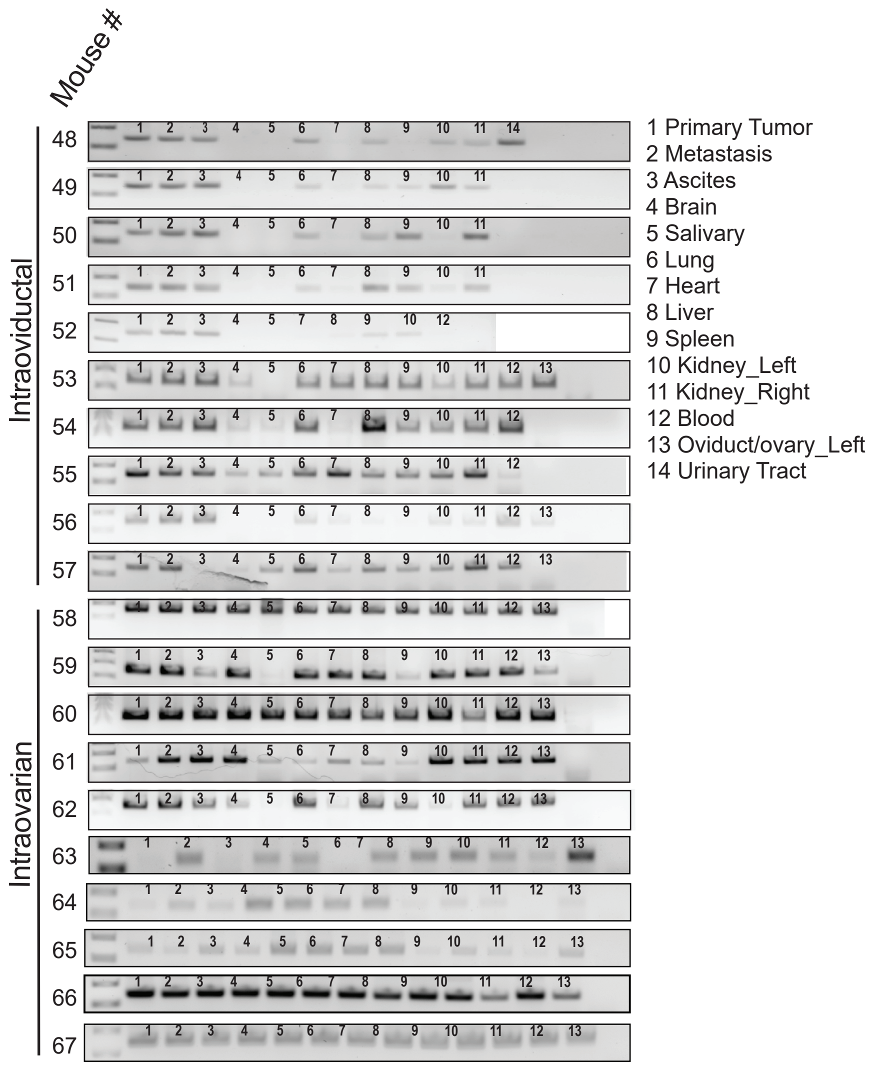

### Figure S8

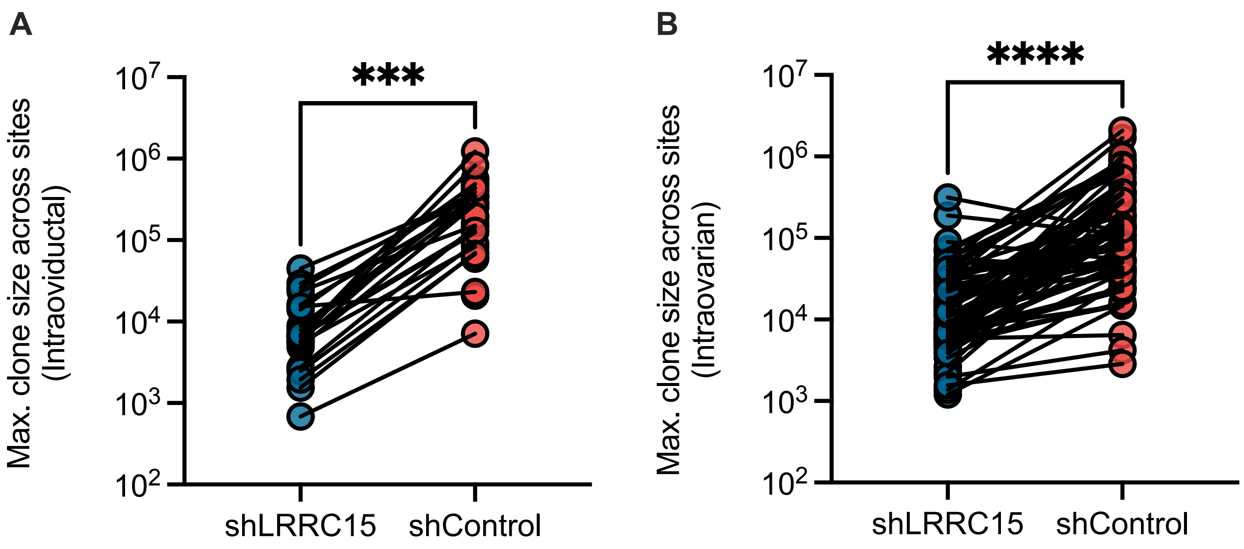

### Figure S9

**Figure S9**

Intraoviductal Injection

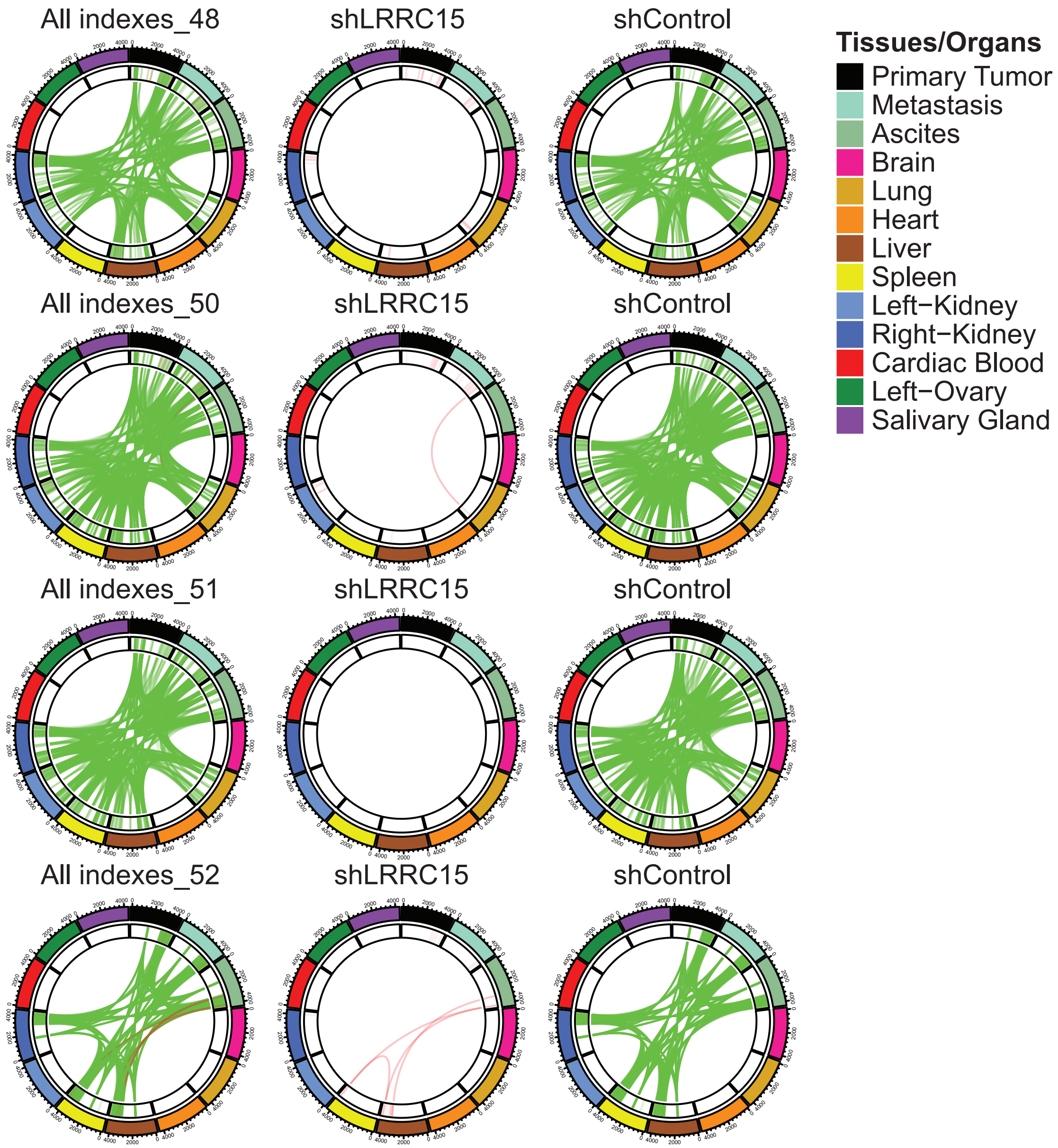

### Figure S10

**Figure S10**

Intraovarian Injection

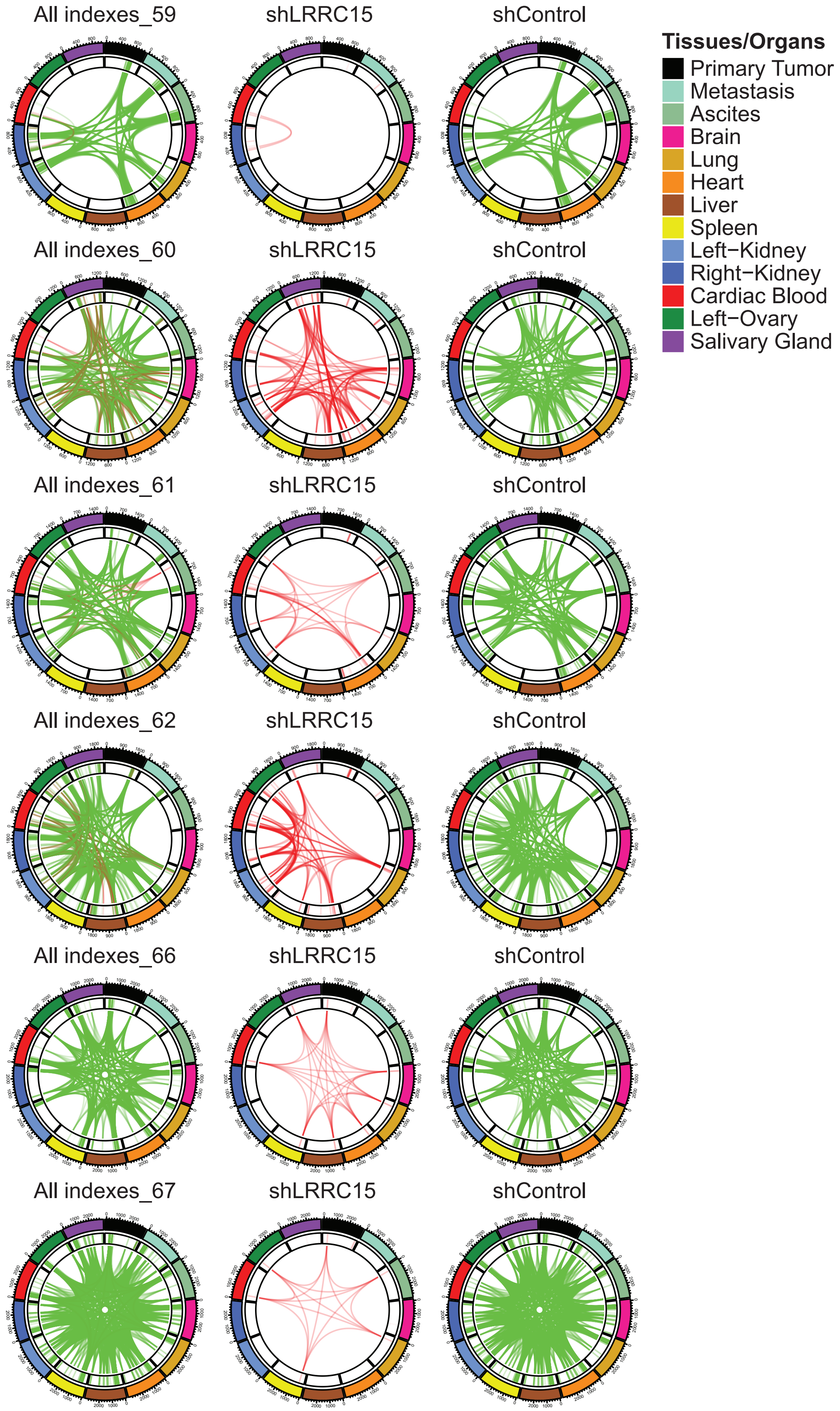

### Figure S11

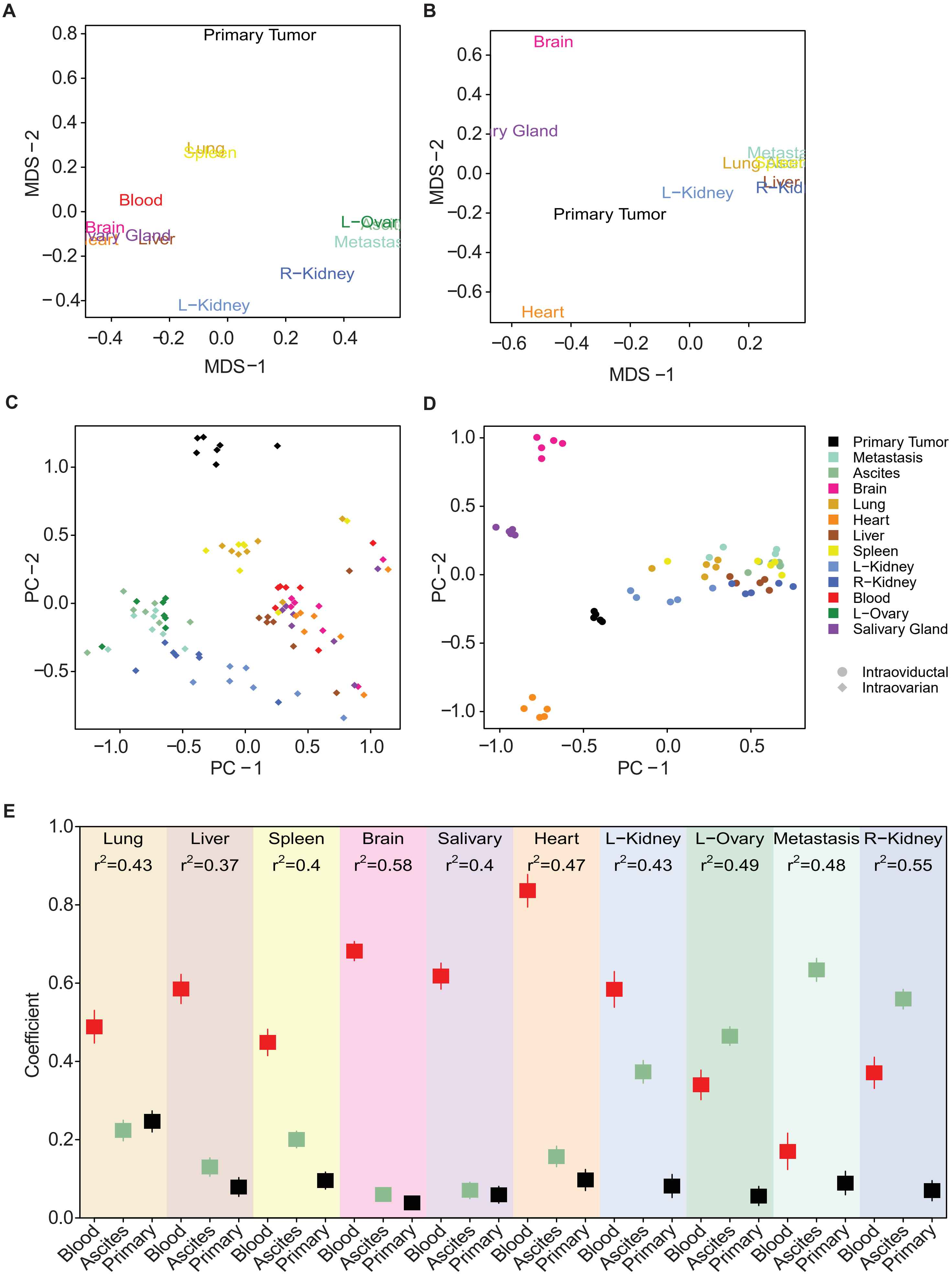

### Figure S12

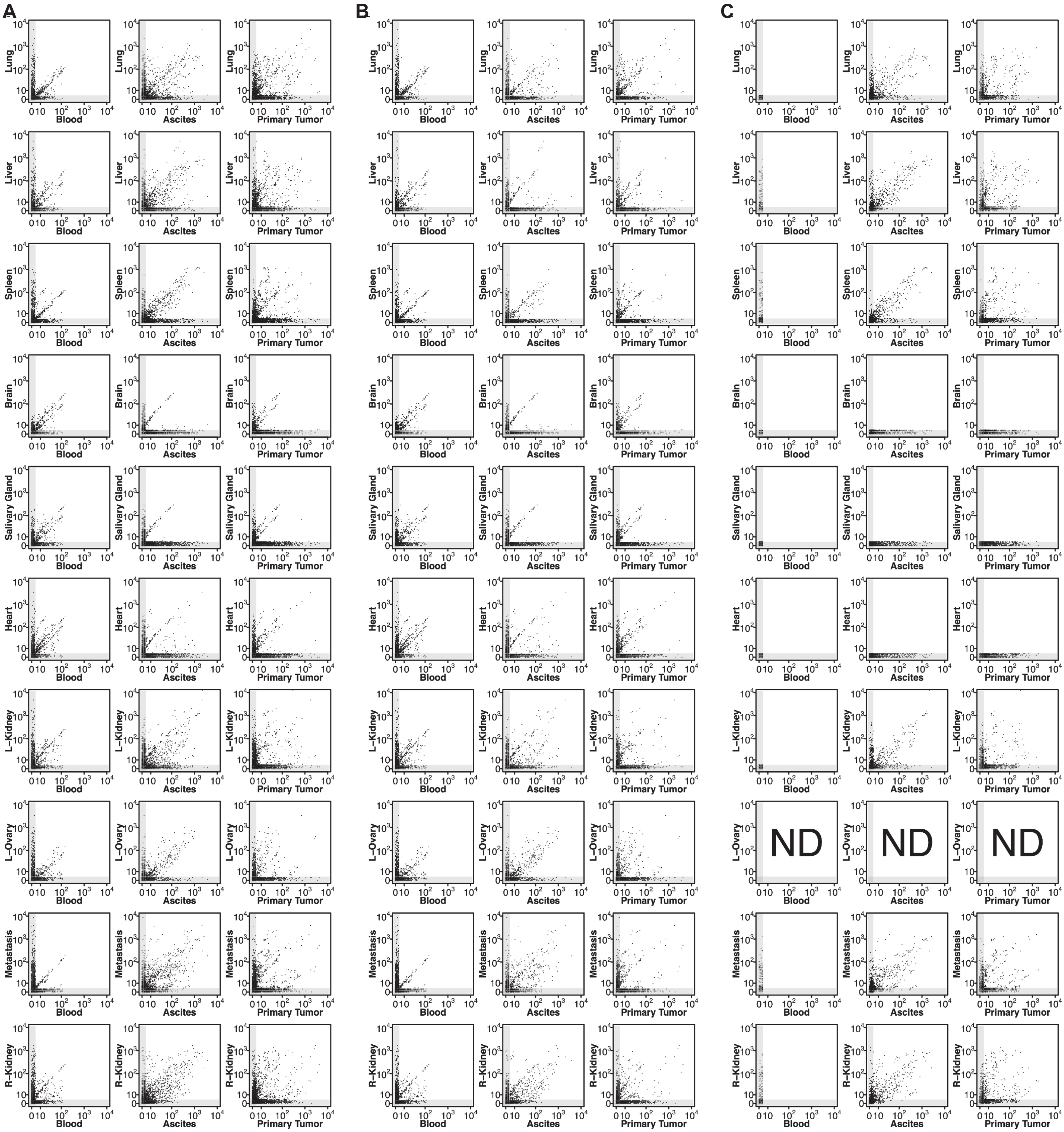

### Figure S13

**Supplementary Figure 13**

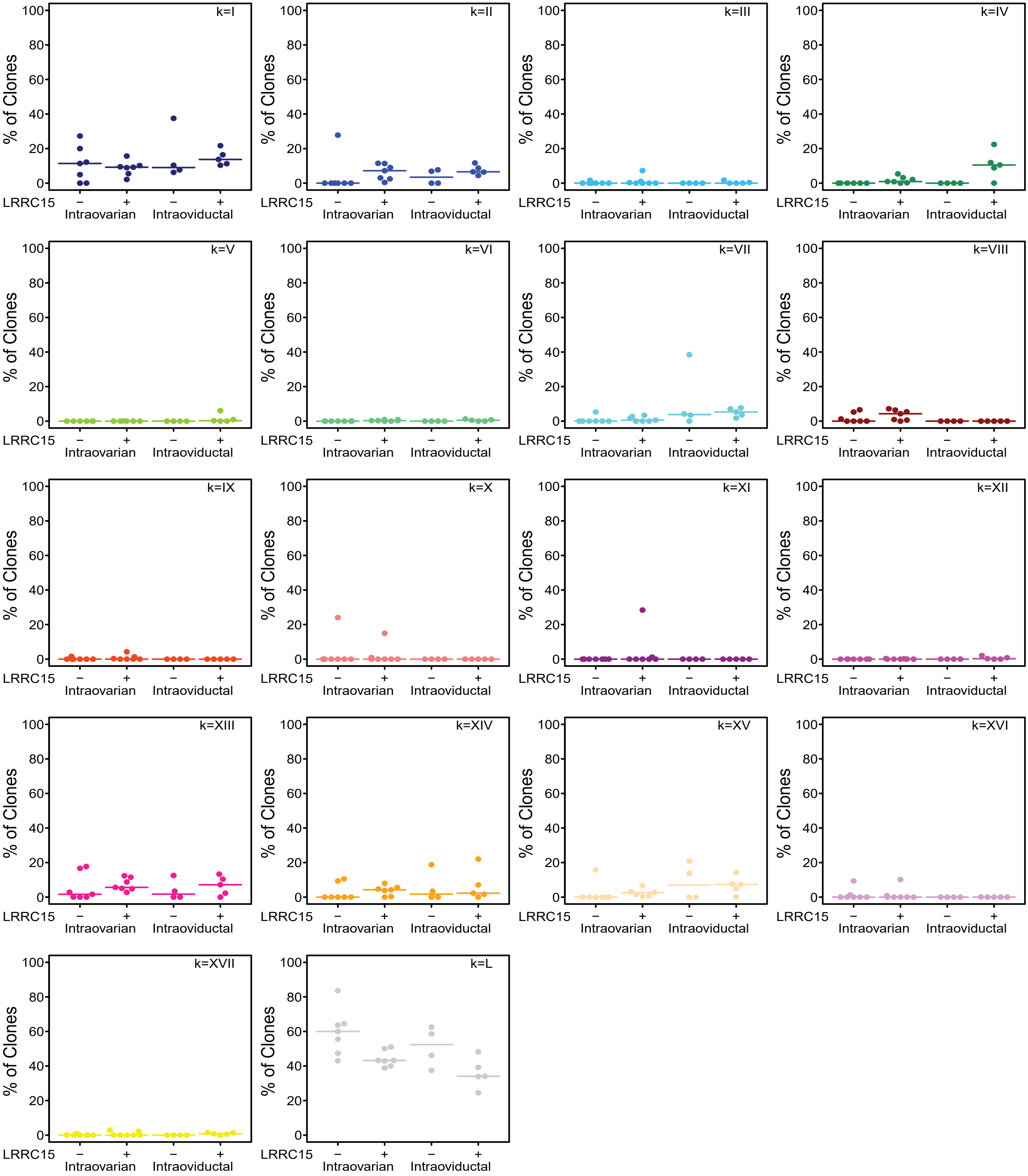

### Figure S14

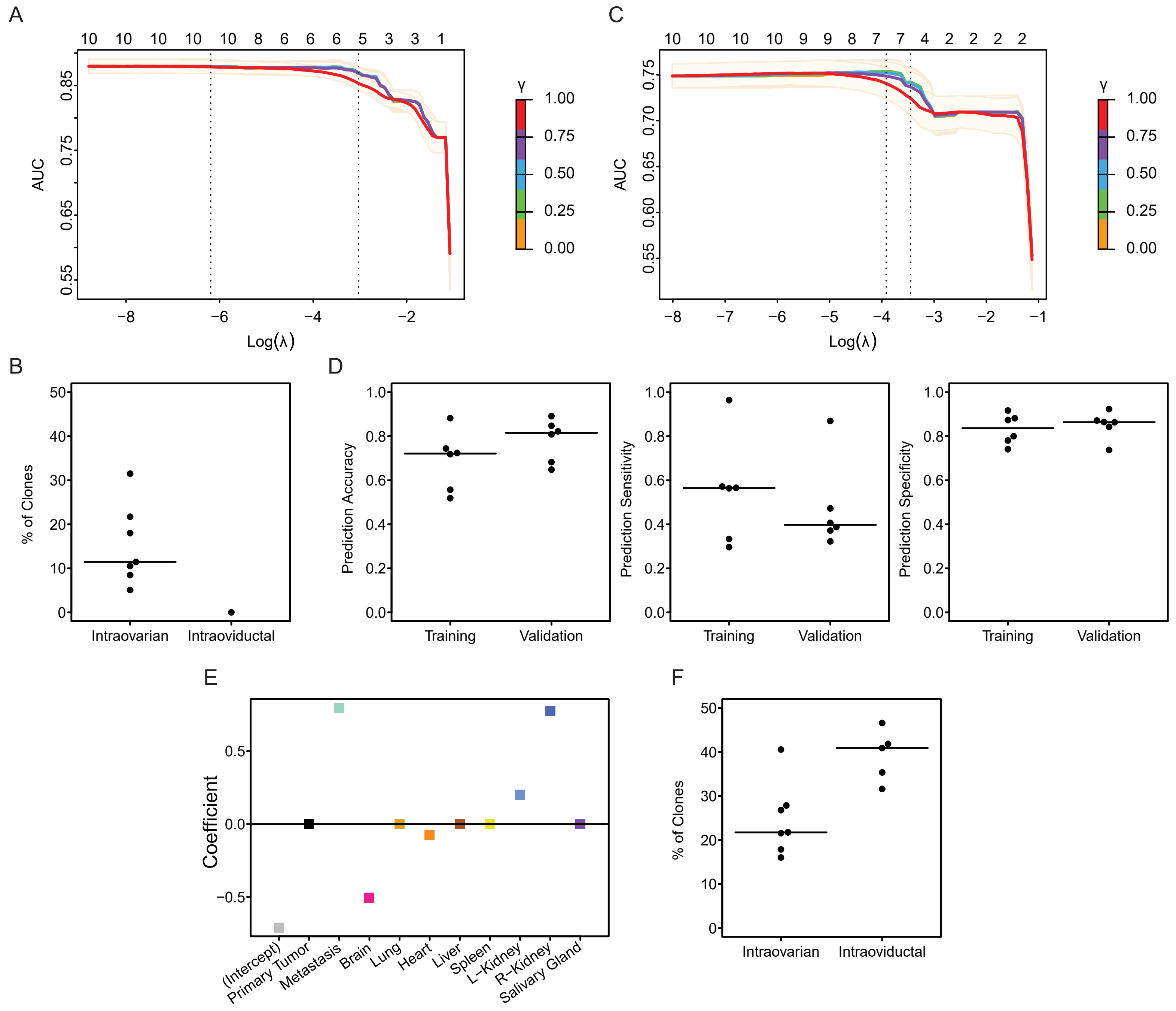
